# An Nrf2/Yap1-like bZIP protein drives UV-induced oxidative stress response in a *Sporobolomyces* yeast with evolutionary conservation

**DOI:** 10.1101/2025.09.17.676916

**Authors:** Ranko Gacesa, Raymond Chung, Suikinai Nobre Santos, Gabriel Padilla, Itamar Soares Melo, Paul F. Long

**Author notes:** Address for correspondence: Professor Paul F Long. Dr. Ranko Gacesa: Department of Gastroenterology & Hepatology, University Medical Centre Groningen, Antonius Deusinglaan, 1, 9713 AV, The Netherlands and Department of Genetics, University Medical Centre Groningen, Antonius Deusinglaan, 1, 9713 AV, The Netherlands. Mr Raymond Chung: Institute of Psychiatry, Psychology and Neuroscience, King’s College London, Denmark Hill, Camberwell, London, United Kingdom. Dr. Suikinai Nobre Santos: Biodiversita Tecnologia Microbiana, Rua Latino Coelho, 1301, Parque Taquaral, 13087-010, Campinas, São Paulo, Brazil.

## Abstract

**Research background:** Ultraviolet radiation represents a major environmental hazard that damages DNA and promotes oxidative stress in all living organisms. In animals, this stress is counteracted by the Nrf2 signalling pathway, while in yeasts such as *Saccharomyces cerevisiae*, the Yap1 transcription factor coordinates oxidative stress responses. The extent to which these mechanisms are conserved in UV-tolerant yeasts from the Basidiomycota lineage remains poorly understood.

**Experimental approach:** We investigated the stress response of a UV-tolerant *Sporobolomyces sp.* isolates exposed to long-term ultraviolet-B irradiation. Yeast survival and antioxidant capacity were assessed, followed by proteome-wide quantification of protein expression using tandem mass tag labelling and multidimensional protein identification technology. Proteins were annotated for biological function through gene ontology and pathway analyses to identify stress-related factors and transcriptional regulators.

**Results and conclusions:** The LEV-2 *Sporobolomyces* yeast isolate displayed remarkable survival under conditions lethal to *S. cerevisiae* and showed induction of antioxidant activity during prolonged exposure. Proteomic profiling revealed large-scale adjustments in protein expression, including the upregulation of enzymes involved in antioxidant biosynthesis, enzymatic antioxidants, DNA repair proteins, heat shock proteins, and multiple signalling pathways. Importantly, several basic leucine zipper proteins with similarity to animal Nrf2 and yeast Yap1 were detected, with one factor showing a distinct induction in response to ultraviolet stress. These findings suggest that the oxidative stress response of *Sporobolomyces* is mediated by conserved regulatory modules comparable to those of both yeasts and animals, highlighting the evolutionary conservation of stress adaptation mechanisms.

**Novelty and scientific contribution:** This work provides the first proteome-level description of the ultraviolet stress response in a Basidiomycota yeast. It demonstrates that bZIP transcription factors drive a complex antioxidant and repair programme in *Sporobolomyces*, functionally reminiscent of the animal Nrf2 pathway. The results support the view that fundamental oxidative stress responses arose early in eukaryotic evolution and have been preserved across distant fungal and animal lineages.

## Introduction

Ultra-violet radiation (UVR) is a major hazard to biological systems and an important environmental source of oxidative stress. The UV-B band of UVR (λ = 280-315 nm) causes direct damage to cellular macromolecules through photo-oxidation [1] and by inducing the formation of DNA photo-adducts [2]. In addition, both UV-A (λ = 315 – 400 nm) and UV-B radiation cause oxidative stress by inducing the production of reactive oxygen species (ROS) in irradiated cells [3], [4]. The basic leucine zipper domain (bZip) transcription factor Nrf2 and its inhibitor, the kelch-domain protein Keap1, play critical roles in regulating the response to oxidative stress in vertebrates [5]. In an unstressed cell, Nrf2 is sequestered in the cytosol by Keap1, ubiquitinated by the Cullin3-Rbx1 E3 ubiquitin ligase – Keap1 (CRLKeap1) complex, followed by proteasomal degradation. During oxidative stress, the increased cellular concentration of ROS causes oxidation of critical “ROS-sensing” cysteine residues of Keap1, which leads to conformational changes in the CRLKeap1 complex and prevents ubiquitination of the Nrf2 protein [6]. The inhibition of CRLKeap1-mediated degradation of Nrf2 results in an increase in cytosolic concentration of newly synthesized Nrf2 and its translocation to the nucleus [7]. Nuclear Nrf2 associates with small Maf proteins and interacts with antioxidant response element (ARE) cis-acting enhancer sequences on DNA to activate transcription of a large number of genes that encode detoxification and antioxidant proteins [8], [9].

In addition to playing a major role in resistance to environmental pollutants [10] and chemically induced oxidative stress [11], the Nrf2 pathway also regulates the response to UV-induced oxidative stress [12]. In a study by Kleszczyński et al., induction of Nrf2-regulated genes by pre-treatment with Nrf2 activators sulforaphane and phenylethyl isothiocyanate was shown to protect human skin cells against UV exposure in cell cultures and ex vivo. In this study, pre-treated, UV-exposed cells had lower rates of apoptosis, reduced levels of sunburn biomarkers, and higher levels of endogenous antioxidants such as catalase when compared to non-treated controls [13]. Nrf2-mediated protection against UV-induced oxidative stress was also demonstrated in mouse models, where treatment with sulforaphane-containing broccoli sprouts significantly reduced the incidence rate of UV-induced skin cancer [14]. Furthermore, a study with genetically modified mouse models found that mouse models lacking an Nrf2-encoding gene recovered slowly from UV-induced inflammation and were highly sensitive to photo-aging [15].

The transcription factor Nrf2 is conserved across vertebrates and invertebrates such as the fly *Drosophila melanogaster* and the worm *Caenorhabditis elegans* [16], but the evolutionary conservation of Keap1-dependent regulation of Nrf2 remains an active area of research. While Keap1-dependent regulation of invertebrate Nrf2 activity was empirically confirmed in *D. melanogaster* [17], the Nrf2 homolog in *C. elegans* did not interact with a kelch-like protein and is instead ubiquitinated by a β-TrCP-ubiquitin ligase complex [18]. This process is similar to Keap1-independent degradation of Nrf2 also observed in vertebrates [19]. A software pipeline was developed for identifying distant homologs of protein and gene sequences, and this software was used to examine databases of animal and microbial genes and proteins for sequences related to human Keap1 and Nrf2 proteins [20]. This data-mining study identified homologs of vertebrate Nrf2 and Keap1 proteins in all publicly accessible animal genomes and in genomes of a large number of fungi, but not in genomes of bacteria or archaea, suggesting that evolution of the Keap1–Nrf2 pathway predated the fungal–metazoan divergence [20]. In a subsequent study, we used fossil records to calibrate the time-frame of emergence of genes encoding Nrf2 in fungi and animals and found that diversification of these genes in animals and fungi occurred simultaneously with the rise in atmospheric oxygen levels during geological time [21]. These studies indicated that bZip transcription factors play a critical part in stress response mechanisms in all eukaryotes, and that the increase in atmospheric oxygen during the Great Oxygenation Event and following geological periods led to evolutionary pressures that drove the evolution of Nrf2-based stress responses in animals [20], [21].

In *Saccharomyces cerevisiae*, UV irradiation activates the RAS/cAMP/PKA signaling pathway independently of DNA damage. This pathway includes the GTPase RAS, which controls the activity of adenylate cyclase (Cyr1) enzyme that stimulates the production of cyclic AMP (cAMP). The increased intracellular concentration of cAMP triggers activation of cAMP-dependent protein kinase A (PKA) and initiates a phosphorylation cascade that activates the bZip transcription factor Gcn4 [22]. Gcn4 is primarily associated with yeast response to starvation and controls transcription of more than 30 genes that encode enzymes involved in amino acid biosynthesis [23]. However, a study of genetically modified *S. cerevisiae* strains by Engelberg et al. (1994) found that *S. cerevisiae* mutants with high, constitutive expression of Gcn4 were ∼3.5-fold more resistant to UV irradiation compared to wild-type yeasts, while strains deficient in the Gcn4-encoding gene were ∼5-fold more sensitive to UV than wild-type yeasts [22]. In addition to RAS signaling, the other major UV-response pathway in *S. cerevisiae* is the Yap1-mediated response to UV-induced oxidative stress [24]. The yeast transcription factor Yap1 is a bZip protein that binds to the AP-1 recognition element in the promoter regions of numerous yeast genes involved in DNA repair and response to oxidative stress, such as glutathione biosynthesis enzymes and thioredoxins [25]. While Yap1 signaling is reminiscent of Nrf2-mediated activation of response to oxidative stress in vertebrates, no interaction between kelch-like proteins and Yap1 has been reported as of yet [26]. In addition to RAS and Yap1 stress response signaling pathways, yeasts also possess DNA damage response pathways. These pathways are largely conserved across all eukaryotes and consist of sensors, primary and secondary signal transducers, and effectors. DNA damage is detected by “sensor” proteins, such as *S. cerevisiae* proteins Rad1, Rad9, and Hus1, and activates signal transduction kinases (Mec1 and Tel1 in *S. cerevisiae*) [27], which activate secondary downstream kinases such as Dun1 [28]. The downstream kinases activate currently unknown effector proteins that increase transcription rates of enzymes involved in DNA repair and also arrest the cell cycle during DNA repair [27].

While *S. cerevisiae* is highly resistant to UV radiation when compared to bacterial models [29], yeasts native to environments with high incidence of solar radiation can tolerate UV radiation levels lethal to *S. cerevisiae*. Examples of such yeasts include carotenoid-containing yeast genera such as *Sporobolomyces* and *Rhodotorula* [29], [30], and the black yeasts of the genus *Exophiala* [29]. The mechanisms of response to UV-induced stress have not been studied in these UV-tolerant yeasts, and it is currently not known whether the stress response signalling mediated by bZip proteins is conserved between UV-tolerant yeasts, *S. cerevisiae*, and the animal Keap1-Nrf2 pathway. UV-tolerant *Sporobolomyces* yeast were selected for this study because of their taxonomic classification (Phylum Basidiomycota as opposed to *S. cerevisiae*, which belongs to Phylum Ascomycota) and to allow study of the effects of high levels of UV-B in the range lethal to *S. cerevisiae*. The aim of this study was to determine, at the proteome level, stress response mechanisms in a *Sporobolomyces* strain designated LEV-2 when exposed to long-term UV-B irradiation, and proteomes of LEV-2 cultures exposed to different durations of UV-B (5 minutes to 24 hours) were quantified using MudPIT mass spectrometry technology. The quantified proteins were functionally annotated using gene ontology (GO) terms [31], KEGG modules, and KEGG pathways [32], [33] to identify proteins involved in stress response and bZip transcription factors. In addition, the fold changes of proteins involved in stress response were examined to describe the stress response of the UV-tolerant yeast LEV-2 over time.

## Materials and methods

### UV-tolerance testing of yeast isolates

*Saccharomyces cerevisiae* and five UV-tolerant yeasts previously isolated and identified by Castelliani et al. [30], designated LEV-2, LEV-9, LEV-12, LEV-13, and LEV-16, were tested for survival after exposure to long-term UV-B irradiation. The yeasts were cultivated in sterile, half-strength YPD liquid medium composed of yeast extract (5 g/L), dextrose (10 g/L), and peptone (10 g/L) in cotton-plugged 250 mL Erlenmeyer flasks at 27 °C with shaking (100 rpm). Three replicates were grown for each sample. After 24 hours of growth, the optical density at 600 nm (OD₆₀₀) of each sample was standardized to OD₆₀₀ = 1.0 (∼3 × 10⁷ cells/mL) by diluting the sample with sterile half-strength YPD liquid medium as necessary, and a 25 mL aliquot of each yeast culture was taken for UV-tolerance testing. The yeast samples were irradiated using dual Philips Ultraviolet-B TL 20W/12RS lamps as follows:

- Sample 1 (Control) was not exposed to UV
- Sample 2 (1h) was irradiated for 1 hour
- Sample 3 (2h) was irradiated for 2 hours
- Sample 4 (4h) was irradiated for 4 hours
- Sample 5 (8h) was irradiated for 8 hours
- Sample 6 (24h) was irradiated for 24 hours

Irradiation was performed at room temperature, and temperature changes during irradiation were assumed to be moderate and not critical for the experimental outcome. During irradiation, each sample was standardized to a volume of 25 mL by adding sterile half-strength YPD medium to account for any evaporation during irradiation. The irradiated samples were vortex-agitated for 30 seconds, and 1 mL aliquots were used to prepare three technical replicates of ten-fold serial dilutions. A 0.1 mL volume of the sample diluted 1:1000, containing ∼3.0 × 10³ cells, was used to inoculate half-strength YPD solid agar. Inoculated petri dishes were incubated at room temperature, and yeast colonies were counted after 48 hours. Survival curves were constructed from the mean values of the replicates, and survival rates were calculated as SR (%) = 100 × [CFU (irradiated sample) / CFU (control sample)].

### Preparation of UV-tolerant yeast isolate for proteomics

The yeast was cultivated in half-strength YPD medium for 24 hours. After 24 hours, cell numbers were estimated at OD₆₀₀ and samples were diluted to OD₆₀₀ = 1.0, equivalent to approximately 3 × 10⁷ cells/mL. Measurements were performed using a UV/VIS spectrophotometer (model 7315, Jenway Ltd., Stone, Staffordshire, UK). The yeast culture was then divided into 20 × 25 mL aliquots (Sample 1 – Sample 10, in duplicate). The samples, in open petri dishes, were irradiated under dual Philips Ultraviolet-B TL 20W/12RS lamps (UVR output of UV-B: 4 J/m²/s and UV-A: 1.75 J/m²/s) at a distance of 10 cm from the lamps. The samples were stirred manually every 20 minutes during irradiation.

Yeast samples were irradiated as follows:

- Sample 1 (Control) was not exposed to UV
- Sample 2 (5’) was irradiated for 5 minutes
- Sample 3 (10’) was irradiated for 10 minutes
- Sample 4 (15’) was irradiated for 15 minutes
- Sample 5 (1h) was irradiated for 1 hour
- Sample 6 (2h) was irradiated for 2 hours
- Sample 7 (4h) was irradiated for 4 hours
- Sample 8 (8h) was irradiated for 8 hours
- Sample 9 (24h) was irradiated for 24 hours
- Sample 10 (Control 2) was not exposed to UV and was kept on the laboratory bench (artificial light, room temperature, no stirring) for 24 hours

During irradiation, each sample was standardized to a volume of 25 mL by adding sterile half-strength YPD medium to account for any evaporation. Samples were vortex-agitated for 30 seconds, and three 1 mL aliquots were taken for DPPH antioxidant assay. The yeast cells in the remaining culture (22 mL) were collected by centrifugation at 1,000 × g for 15 minutes at room temperature. The supernatant was discarded, and the cells were transferred to 1.5 mL microcentrifuge tubes. To remove any residual liquid medium, pellets were re-suspended in phosphate-buffered saline (PBS) buffer and the cells were collected by centrifugation at 12,300 × g for 15 minutes at room temperature. This procedure was repeated twice. Pellets were flash-frozen by immersion in liquid nitrogen and stored at -80 °C until proteomics analysis; frozen samples were transported on dry ice.

### DPPH assay of extracts from isolated yeast cultures

Yeast cells were collected by centrifugation (5 minutes, 15,000 × g, room temperature) and re-suspended in 1 mL of cell lysis buffer composed of 50 mM tris-buffered saline (TBS) pH 7.6, mixed with 0.1% (w/v) Triton X (Sigma Aldrich). The cells were disrupted using a sonicator probe (Model: VC250, Sonics & Materials Inc.) with a duty cycle of 40% and an output of 3. Samples were kept on ice, and sonication was performed in 10 × 1-minute cycles with 1-minute pauses between cycles. Cell lysis was confirmed by examination of the samples using a light microscope at ×40 magnification. Cell debris was removed by centrifugation (5 minutes, 15,000 × g, room temperature), and the supernatant was tested for free radical-quenching antioxidant activity using a colorimetric assay based on the neutralization of stable free radical 2,2-diphenyl-1-picrylhydrazyl (DPPH) [34]. Briefly, 0.1 mL of each sample was mixed with 1.5 mL of 70 μM DPPH dissolved in methanol. The samples were shielded from light with aluminum foil and incubated for 30 minutes at room temperature, and the color change from violet to yellow, which occurs when DPPH is reduced upon reaction with an antioxidant, was recorded at 515 nm using a UV/VIS spectrophotometer (model 7315, Jenway Ltd., Stone, Staffordshire, UK). A mixture of half-strength YPD medium (0.1 mL) and DPPH (1.5 mL) served as a control, and a mixture of methanol (0.1 mL) and DPPH (1.5 mL) served as the reaction blank. The percentage of DPPH radical scavenging activity was calculated as: Scavenging activity (%) = 100 × (A_blank - A_sample)/A_blank. Experiments were performed in technical triplicates with three replicates, and scavenging activities were plotted as the mean of the 9 triplicate/replicate values against compound concentration.

### Mass spectrometry analysis

The protein composition of the yeast samples was identified using Multidimensional Protein Identification Technology (MudPIT), with Tandem Mass Tags (TMT) used for relative quantification of labelled peptides, as follows: Ten frozen yeast cell pellets (Sample 1 – 10, in duplicate) were processed for proteomics analysis. Cell pellets were homogenized in 100 μL of ice-cold lysis buffer using a micro-pestle. The lysis buffer consisted of 100 mM Tris(hydroxymethyl)aminomethane hydrochloride (pH 7.5), 300 mM NaCl, 2% (v/v) Triton X-100, 1% (w/v) sodium deoxycholate, 200 mM NaOH, 2% (w/w) SDS, 2% (v/v) β-mercaptoethanol, 2× Complete Protease Inhibitor Cocktail (Roche), and 2× Complete Phosphatase Inhibitor Cocktail (Roche). The cell pellets were homogenized for 30 seconds and placed on ice for 1 minute. This procedure was repeated four more times. Cell debris was pelleted by centrifugation (14,000 × g, 14 minutes, 4 °C) and the supernatants containing solubilized proteins were transferred to fresh 1.5 mL tubes and kept on ice. For each yeast sample, the solubilized proteins were transferred into a fresh 2 mL microcentrifuge tube for protein precipitation. The solubilized proteins were combined with 800 μL of methanol, vortex-agitated for 30 seconds, and centrifuged at 14,000 × g for 1 minute at room temperature. The sample was mixed with 200 μL of chloroform, vortex-agitated for 30 seconds, and centrifuged at 14,000 × g for 1 minute at room temperature. Six hundred microliters of deionized water was added to the mixture, and the mixture was vortex-agitated for 30 seconds and centrifuged at 14,000 × g for 10 minutes at room temperature. The upper aqueous phase of the sample was discarded, 800 μL of methanol was added to the remainder of the sample, and the precipitated proteins were collected by centrifugation at 14,000 × g for 5 minutes at room temperature. The supernatant was discarded and the pellet consisting of precipitated proteins was dried in a Savant SpeedVac Concentrator (Thermo Fisher Scientific) for 1 minute at room temperature. Precipitated proteins were solubilized in 150 μL of buffer composed of 100 mM Triethylammonium bicarbonate (TEAB) and 10 mM Tris(2-carboxyethyl)phosphine hydrochloride (TCEP) (Sigma). Protein pellets were dissolved by 10 minutes of vortex agitation at room temperature. A 10 μL aliquot was taken for protein concentration determination by Bradford assay (Bio-Rad) using bovine serum albumin (Sigma) as a standard, and the remaining sample was incubated in the buffer (composed of 100 mM TEAB and 10 mM TCEP) for 1 hour at 55 °C to reduce protein disulfide bonds. For each yeast sample, a volume containing 30 μg of proteins (as calculated from Bradford assay) was transferred to a 1.5 mL tube and adjusted to 100 μL with buffer containing 100 mM TEAB and 10 mM TCEP (Sigma). Samples were alkylated by addition of 10 μL of buffer containing 100 mM TEAB, 10 mM TCEP, and 198 mM iodoacetamide (Sigma) and incubated for 30 minutes at room temperature. The incubating samples were protected from light with aluminium foil. Proteins were digested by adding 10 μL of trypsin (Promega) solution (60 ng/μL) in 100 mM TEAB and incubating overnight at 37 °C, protected from light with aluminium foil.

Tandem Mass Tag (TMT) 10-plex reagents (Thermo Fisher Scientific TMT 10-plex kit, https://www.thermofisher.com/order/catalog/product/90110) were reconstituted according to the manufacturer’s instructions by adding 41 μL of acetonitrile (ACN) to 0.8 mg of each TMT label. The appropriate reconstituted label was added to the protein digests of yeast samples as follows:

| Sample | TMT Label | Sample | TMT Label |
| --- | --- | --- | --- |
|  | Reagent |  | Reagent |
| **1 (Control)** | 126 | 6 (2 hours of UV) | 129N |
| **2 (5 min of UV)** | 127N | 7 (4 hours of UV) | 129C |
| **3 (10 min of UV)** | 127C | 8 (8 hours of UV) | 130N |
| **4 (15 min of UV)** | 128N | 9 (24 hours of UV) | 130C |
| **5 (1 hour of UV)** | 128C | 10 (Control) | 131 |

Samples were vortex-agitated for 30 seconds and incubated at room temperature for one hour. The labelling reactions were quenched by the addition of 9 μL of 5% (v/v) hydroxylamine (Sigma) and incubated for 15 minutes. The 10 samples were combined in equal amounts of 10 μL. Excess labelling reagent was removed by solid-phase extraction using a 1 mL Oasis HLB cartridge (Waters) as follows: The sample was prepared for purification in 4% (v/v) ACN and 0.1% (v/v) trifluoroacetic acid (TFA). A vacuum manifold was used to apply buffers and sample to the solid-phase extraction cartridge at a rate of 1 mL/minute. The solid-phase extraction cartridge was conditioned with 2 × 1 mL of conditioning buffer consisting of 95% (v/v) ACN and 0.1% (v/v) TFA in deionized water. The conditioned cartridge was washed twice with 1 mL of wash buffer composed of 5% (v/v) ACN and 0.1% (v/v) TFA in deionized water. The sample was applied to the cartridge; the flow-through was collected and re-applied to the cartridge. Unbound contaminants were washed through with 5 × 1 mL of wash buffer composed of 5% (v/v) ACN and 0.1% (v/v) TFA in deionized water, and the bound labeled peptides were eluted with 3 × 1 mL of elution buffer composed of 85% (v/v) ACN and 0.1% (v/v) TFA in deionized water. The eluted labelled peptides were lyophilized in a SpeedVac (2 hours at room temperature).

The TMT-labelled samples 1 - 10, containing lyophilized peptides labelled with TMT reagents, were reconstituted in 1.8 mL OFFGEL buffer consisting of 9.6% (v/v) glycerol and 0.96% (v/v) ampholytes in the form of IPG buffer pH 3-10 (GE Healthcare Life Sciences) in deionized water. The peptides were solubilized using a sonic water bath for 10 seconds, followed by 30 minutes of vortex agitation at room temperature, and insoluble material was removed by 15 minutes of centrifugation at 14,000 × g at room temperature. The supernatant was applied to an isoelectric focusing (IEF) strip pH 3-10 (GE Healthcare Life Sciences) according to the manufacturer’s instructions, and the labeled peptides were separated into 12 fractions and collected using an Agilent 3100 OFFGEL Fractionator. Isoelectric focusing was performed at 20 °C for a total of 20 kVh at a constant current of 50 μA. Once completed, fractionation was held at 500 V until fraction collection. The fractions were collected in 1.5 mL tubes and acidified by the addition of TFA (final acid concentration of 0.1% [v/v]). For each fraction, salts, TFA, and gel debris were removed by solid-phase extraction using the procedure described above, with the exception that the fractionated peptides were eluted in 1 mL of buffer composed of 85% (v/v) ACN and 0.1% (v/v) formic acid in deionized water. Eluted peptides were lyophilized in a SpeedVac (2 hours at room temperature).

The peptides were solubilized in 50 mM ammonium bicarbonate for separation of the peptide mixture by liquid chromatography and analysis by tandem mass spectrometry. A portion of each fraction was analysed sequentially from fraction 1 (pH 3) to fraction 12 (pH 10). Chromatographic separations were performed using the Ultra-High Performance Liquid Chromatography (UHPLC) system EASY-nLC II (Thermo Fisher Scientific). The peptides were separated using a reverse-phase chromatography column 100 mm EASY-Column with an internal diameter of 75 μm packed with a stationary phase of C18, 3 μm particles, and 120 Å porosity (Thermo Scientific). Peptides were eluted using a gradient of ACN (5% to 40% over 100 minutes, increased to 80% over 10 minutes and held at 80% for 5 minutes) and 0.1% (v/v) formic acid. The flow rate of solvent was 300 nL/minute. Mass spectra were acquired on the LTQ Orbitrap Velos Pro (Thermo Fisher Scientific) operated by Xcalibur™ software. The instrument was set to record mass spectra ranging from 350 to 1800 m/z at a resolution of 30,000. The 10 most intense precursor ions were subjected to sequencing by high-energy collision-induced dissociation (CID) in the ion trap with a threshold of 5000 counts. The precursor ion selection isolation width was 2 units, and the normalized CID energy for precursor ion fragmentation was 35. Automatic gain control settings for FTMS survey scans were 10⁵ counts and FT-MS/MS scans were 10³ counts. Maximum acquisition time was 500 ms for survey scans and 250 ms for MS/MS scans. Charge-unassigned and single-charge state ions were excluded from MS/MS analysis.

### Data analysis: database searching

A database for spectra matching was constructed from the following sources:

- UniProt (www.uniprot.org) yeast protein sequences for *Sporidiobolus salmonicolor* and *Rhodosporidium toruloides*.
- *Rhodotorula minuta* proteome (designation Rhomi1) acquired from The Fungal Genomics Resource (http://genome.jgi.doe.gov/programs/fungi/index.jsf).
- *Sporobolomyces linderae* CBS 7893 proteome (designation Spoli1) acquired from The Fungal Genomics Resource (http://genome.jgi.doe.gov/programs/fungi/index.jsf).
- *Sporobolomyces roseus* proteome (designation Sporo1) acquired from The Fungal Genomics Resource (http://genome.jgi.doe.gov/programs/fungi/index.jsf).
- Fungal basic leucine zipper sequences assembled using BLASTp [35] search of the NCBI non-redundant (NR) database [36].

Due to limitations of Mascot software, which does not support merging of databases that utilize different formats of protein sequence identifiers (such as UniProt and GenBank), each dataset was processed independently, and the results were merged after database matching of tandem mass spectra. Database matching of MS/MS spectra was performed using Mascot software, version 2.2.03 (Matrix Science). Databases were installed in Mascot; Xcalibur raw files were processed into peak lists with Proteome Discoverer 1.4 (Thermo Fisher Scientific). The Proteome Discoverer Daemon was used to process the raw files with multidimensional protein identification technology (MudPIT) specifying up to 3 missed cleavages, a precursor ion mass tolerance of 20 ppm, and a fragment ion tolerance of 0.8 Da. A variable/dynamic modification for oxidized methionine was set. Fixed/static modifications for carbamidomethylated cysteine and TMT-tagged lysine and N-termini were set. A target false discovery rate (FDR) for high confidence peptide hits was set to 0.01 (1%), and a target FDR for medium confidence peptide hits was set to 0.05 (5%). An independent search was conducted to assess labeling efficiency by specifying all modifications as variable/dynamic. For all high and medium confidence peptides, 98% were modified by TMT, 95% of these were N-terminally labeled, and 96% of lysines were modified by TMT labels.

### Data analysis: quantification and result pre-processing

Detected proteins were grouped under the strict maximum parsimony principle. All detected peptides with a TMT modification of the N-terminus were used to determine a normalization factor for each label. All available reporter ion intensities were summed for each individual label, and the median of the summed intensities was determined for the ten labels. The normalization factor for each label was obtained by expressing the median intensity over the sum of intensities and applied to the raw reporter ion intensities for each respective label. Peptides for which all reported TMT reporter ions were detected and quantified were retained for further analysis. For each protein with multiple quantified peptide hits, the protein signal intensity was calculated as the mean value of TMT reporter ion intensities of peptides matched to the protein. The expression fold-changes of samples were calculated relative to Sample 1 (non-irradiated control). All quantified proteins were annotated using InterProScan software [37] to predict the likely biological functions based on Gene Ontology (GO) terms [31] and Pfam profiles [38]. Protein sequences were also annotated using BlastKOALA software [32] to assign KEGG pathways and KEGG modules [33]. The results of computational annotation were manually curated by examining the primary literature and the information deposited in UniProt and NCBI protein databases. Proteins for which computational annotation was not successful or resulted in prediction of “predicted protein” or “unknown protein” were manually annotated by examining the results of BLASTp searches of UniProt and NCBI SwissProt databases with an E-value cut-off of 0.001, and function was assigned to the protein based on related proteins if possible. Computational annotations and visualization of results were conducted using in-house R scripts and Python scripts.

### Statistical analysis of annotated proteins with increased fold changes after UV-B exposure of yeast cells

Statistical analysis used Fisher’s exact test [39] to determine the GO terms over-represented (more frequent than would be expected based on random distribution) in the dataset of proteins with significant (2-fold or greater) fold change in at least one UV-exposed sample. The test was conducted using the following procedure:

1. Quantified proteins were separated into sensitive dataset (D_s) and control dataset (D_c). Proteins were included in D_s if protein fold changes in at least one UV-exposed yeast sample were 2.0 or higher. The remaining proteins were included in D_c.
2. For each GO term (tested GO term) from the list of all GO terms assigned to quantified proteins, the number of proteins was determined for: a) Proteins in D_s annotated with the tested GO term (n_s^GO) b) Proteins in D_c annotated with the tested GO term (n_c^GO) c) Proteins in D_s annotated with other GO terms (n_s^O) d) Proteins in D_c annotated with other GO terms (n_c^O)
3. One-sided Fisher’s exact test for over-representation of tested GO term in D_s was performed on the following frequency distribution table:

|  | Tested GO | Other GO |
| --- | --- | --- |
| Proteins in $D_s$ | $n_s^{GO}$ | $n_s^O$ |
| Proteins in $D_c$ | $n_c^{GO}$ | $n_c^O$ |

The null hypothesis (H₀) of the test was that the tested GO term is equally frequent amongst the proteins in dataset D_c and amongst the proteins in dataset D_s, while the alternative hypothesis (H_A) was that the tested GO is over-represented amongst the proteins from D_s. The tested GO term was considered significantly over-represented for Fisher’s exact test p-value ≤ 0.05. To assess the robustness of this approach, equivalent Fisher’s exact tests were also performed where proteins were included in the D_s dataset if the protein fold-changes were 1.5 or higher and if the fold-changes were 4.0 or higher. In the results presented, the GO terms were considered significantly over-represented if p-values of at least 2 out of these 3 tests were below the 0.05 threshold, or if the p-value was below 0.05 in the fold-change ≥ 4.0 test. Statistical analysis of over-representation of KEGG Pathways and KEGG Modules was performed using the methodology for statistical analysis of GO terms (described previously), with the exception that the GO terms were replaced with KEGG Pathway terms or KEGG Modules during step 2) of the statistical analysis.

### Statistical analysis of annotated proteins with fold change reduction after UV-B exposure of yeast cells

The statistical analysis was conducted to determine the GO terms, KEGG pathways, and KEGG modules significantly over-represented in the dataset of proteins showing fold change reduction in UV-B exposed yeast cultures. This analysis was conducted using the methodology described in the previous section, with the exception that proteins were included in the sensitive dataset (D_s) if the proteins showed fold change reduction of 2.0 or higher in at least one UV-exposed yeast culture.

## Results

### Determining the UV tolerance of the yeast samples

Five yeast samples previously isolated by Castelliani et al. (LEV-2, LEV-9, LEV-12, LEV-13, and LEV-16) [30], along with *S. cerevisiae*, were tested for viability after exposure to UV-B irradiation. *S. cerevisiae* was used as a control based on the study by Pulschen et al. (2015), who used it as a UV-sensitive yeast control to evaluate the viability of UV-resistant yeasts from high-altitude extreme environments [29]. The survival rates of yeasts exposed to UV-B irradiation are shown in Figure 1. The UV tolerance of these yeasts was no higher than the *S. cerevisiae* control, and these isolates were not considered for further study. The yeasts LEV-2, LEV-13, and LEV-9, however, showed high survival rates after 24-hour exposure to UV-B and were next evaluated for antioxidant activity. Extracts of these yeasts, prepared by cell lysis and removal of insoluble material, were tested using a colorimetric assay based on the quenching of the stable free radical 2,2-diphenyl-1-picrylhydrazyl (DPPH) (Figure 2). Extracts of all three tested yeasts showed an increase in DPPH quenching activity after 24 hours of UV-B irradiation. The increase in antioxidant activity was moderate (∼25% increase) for samples LEV-9 and LEV-13, while the extract of yeast LEV-2 showed a ∼75% increase in antioxidant activity after 24 hours of UV-B irradiation. Yeast LEV-2 also showed a ∼50% reduction in DPPH quenching for samples exposed to 1 hour and 2 hours of UV-B.

**Figure 1:**
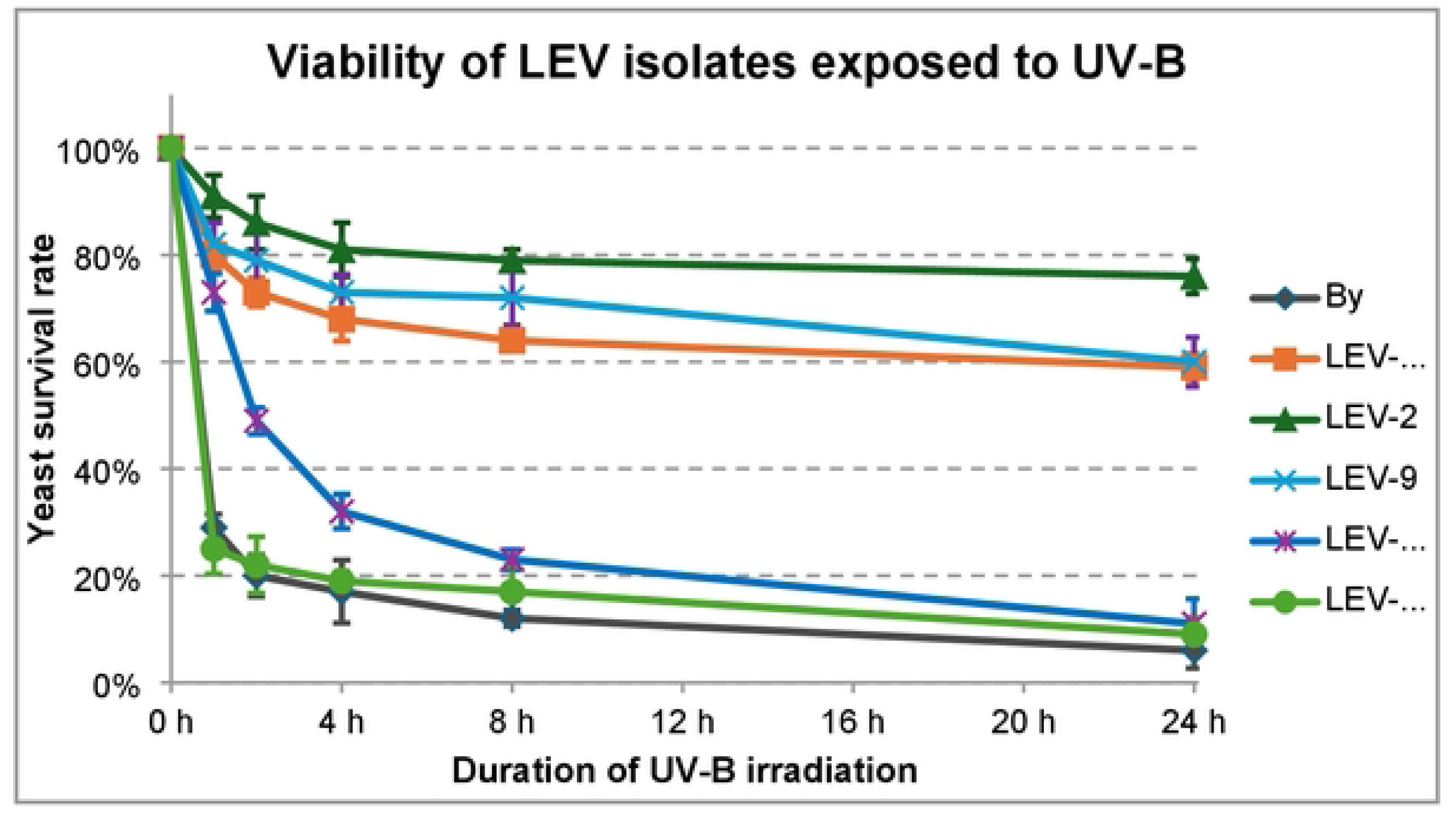
Survival curves of LEV yeast isolates upon exposure to UV-8. UV-B exposure for 24 hours was lethal to S. *cerevisiae,* while the yeasts LEV-12 and LEV-16 showed survival rates below 10 %. The isolates LEV-2, LEV-9 and LEV-13 showed survival rates above 60 %. The isolate LEV-2 had the highest rate of survival amongst the tested yeasts with a survival rate of -80 % after 8 hours of UV-B exposure and -75 %, after 24 hours of UV-B exposure.

**Figure 2:**
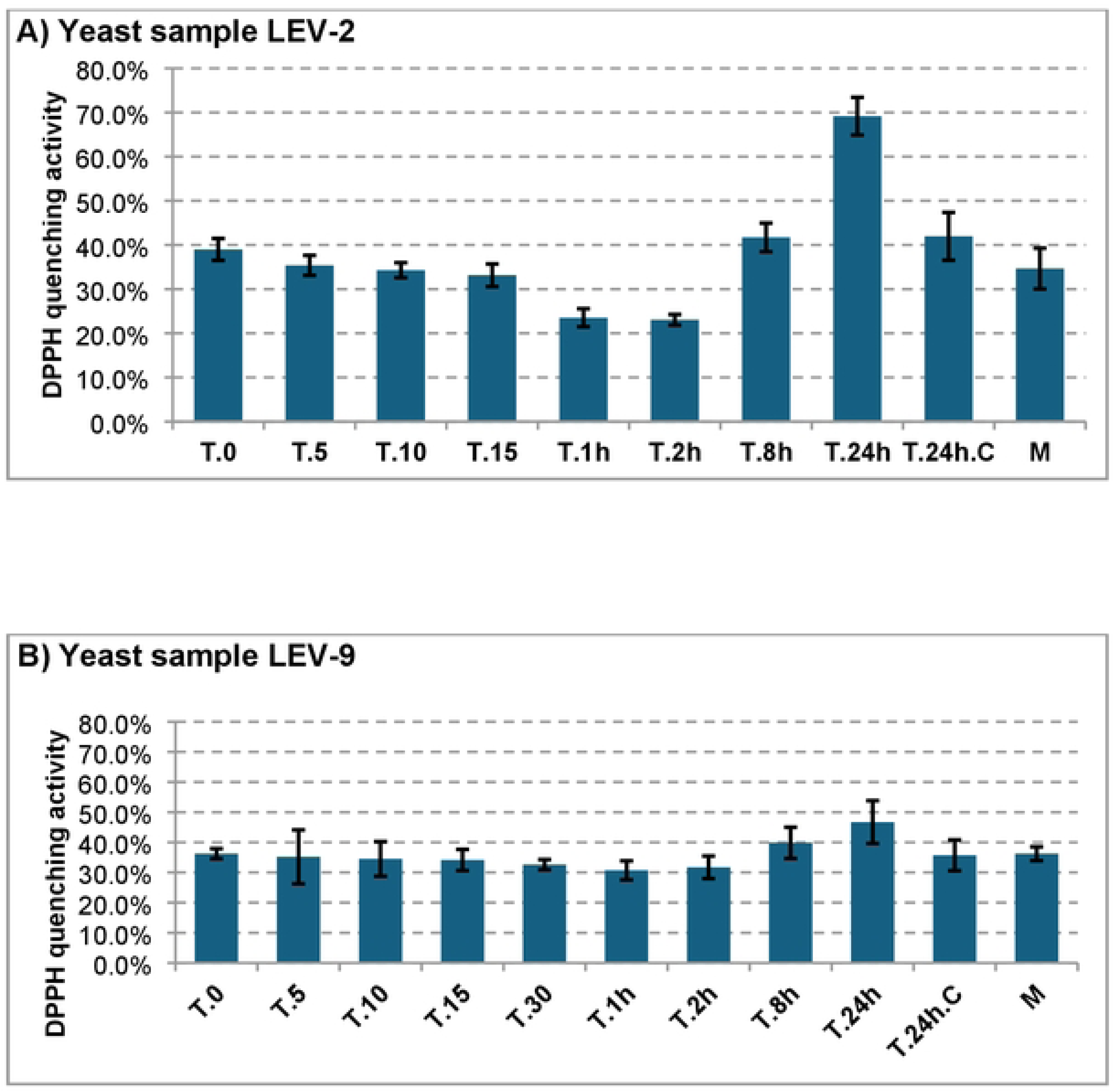
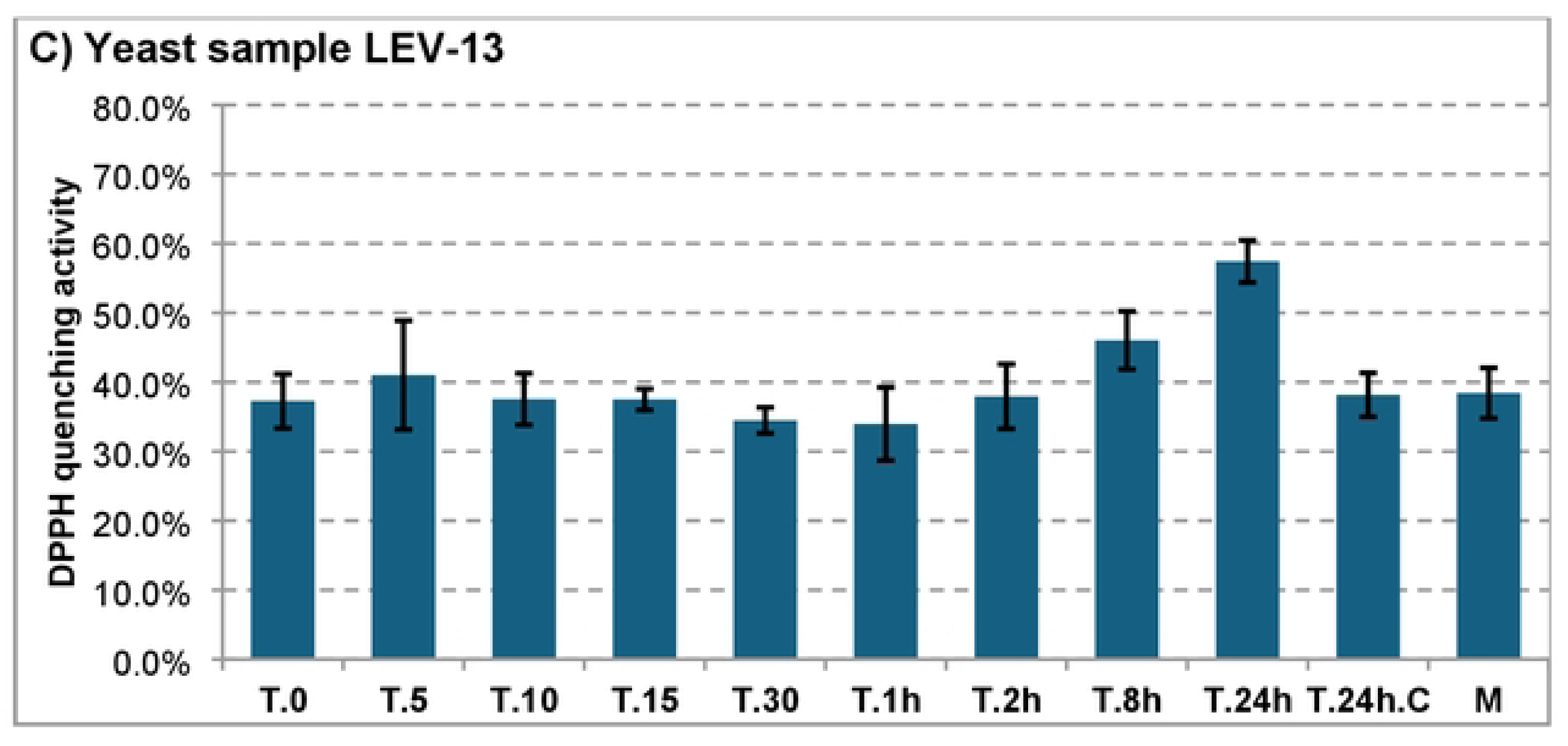
DPPH quenching activity of LEV yeast isolates exposed to UV-B. The figure displays the DPPH free radical quenching activity of extracts of yeast isolates LEV-2 (A), LEV-9 (8) and LEV-13 (C). Yeast cultures were irradiated for time periods ranging from 5 minutes (sample T.5) to 24 hours (sample T.24), using dual Philips Ultraviolet-8 TL 20W/12RS lamps, and yeast extracts were obtained by cell lysis and removal of insoluble material. The controls were half-strength YPD medium (sample M}, non-irradiated yeast cultures (sample T.O), and non-irradiated yeast cultures grown for 24 hours (sample T.24.C). Error bars indicate 1 standard deviation of the mean, calculated from three experiments; DPPH quenching values of three technical triplicates of each experiment were averaged.

The extract of yeast LEV-2, previously identified as *Sporobolomyces* sp. [30], was chosen as the model organism for further study because of its high tolerance to UV-B (Figure 1) and the observed induction of DPPH free radical quenching activity (Figure 2). To assess whether this yeast adapts to UV-induced stress during UV-B exposure, the rate of yeast cell death was calculated for different time periods of UV-B exposure. The results presented in Figure 3 indicated that the initial two hours of UV-B exposure caused moderate loss of viability in yeast LEV-2 (∼9% during the first hour, followed by ∼6% reduction in viability in remaining yeasts in the second hour), but the rate of cell death diminished over the following time periods, and the cell death rate of LEV-2 yeast culture was ∼0.6% of viable CFU per hour after 8 hours of UV-B exposure.

**Figure 3.**
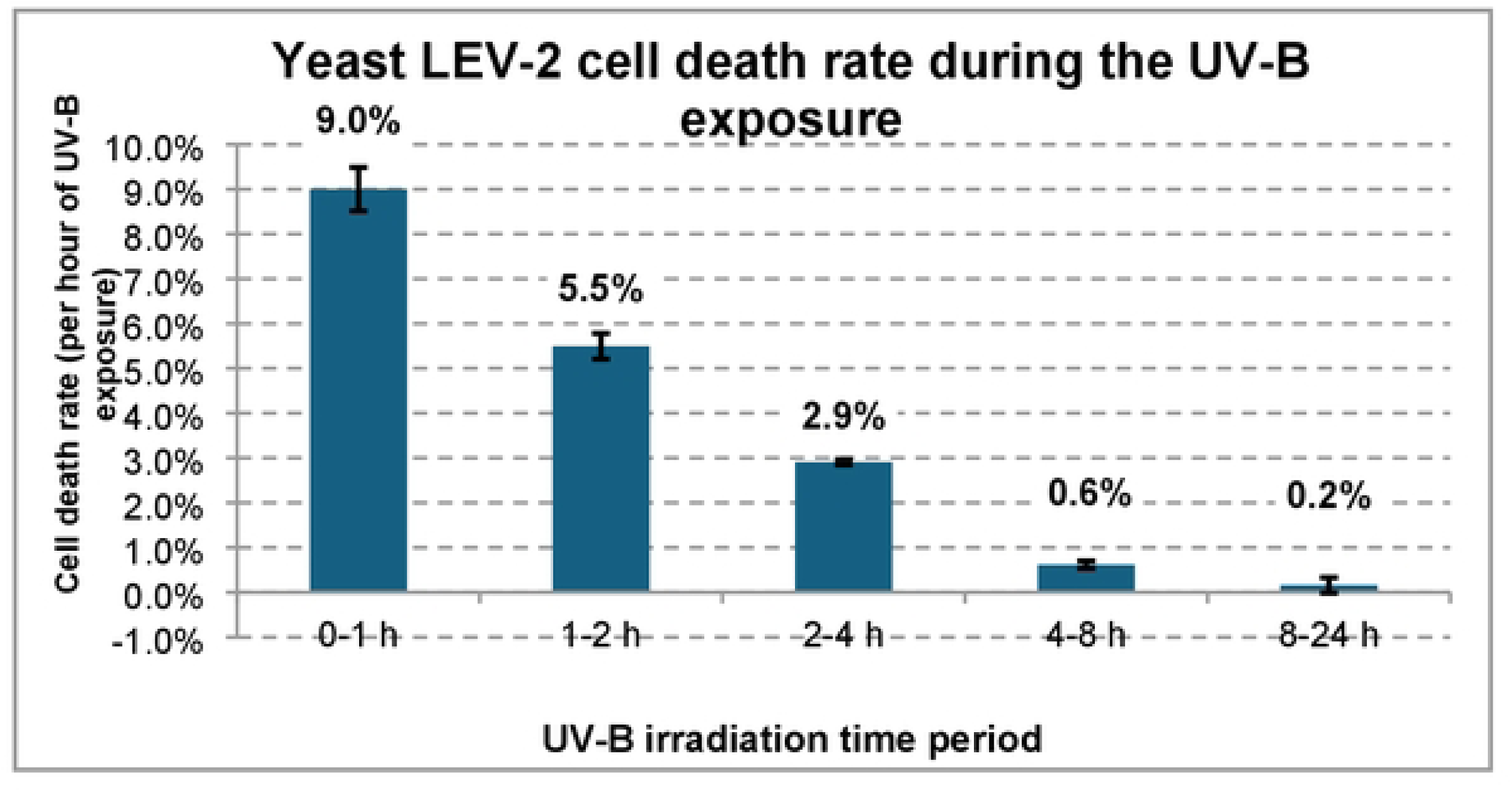
Death rate of yeast LEV-2 exposed to UV-B radiation. The figure shows the rate of cell death of LEV-2 yeast irradiated using dual Philips Ultraviolet-8 TL 20W/12RS lamps for each time period. The cell death rate values are denoted above bars and are expressed as percentage reduction in number of viable colony forming units per hour of UV-8 irradiation. The error bars represent one standard deviation of the mean of three experimental replicates.

### MudPIT analysis of UV-tolerant yeast LEV-2

*Sporobolomyces* yeast LEV-2 was grown in liquid medium and exposed to increasing durations of UV radiation. Irradiation was performed for time intervals ranging from 5 minutes to 24 hours, with UV output of 4 J/m²/s UV-B and 1.75 J/m²/s UV-A. Solubilized proteins were extracted from UV-irradiated samples and non-irradiated controls, labelled with TMT chemical tags, and quantified using multidimensional protein identification technology (MudPIT). The analysis identified 751 proteins for which fold changes could be determined (data not shown).

Based on previously published proteomics studies [40–42], protein expression fold changes of 2 or higher were considered significant. A total of 227 proteins (∼30% of quantified proteins) showed a significant fold change increase (2 or higher) in irradiated yeast LEV-2 cultures (Figure 4A). The median value of fold changes of these 227 proteins was ∼1 for LEV-2 controls and for samples exposed to UV-B for up to 4 hours; ∼1.5 for LEV-2 exposed to 8 hours of UV-B; and ∼2.5 for LEV-2 exposed to 24 hours of UV-B. A total of 279 proteins (∼37% of quantified proteins) showed significant fold change decrease in UV-irradiated yeast cultures (Figure 4B). For these 279 proteins, the median fold change value was ∼1.0 for non-irradiated controls and for LEV-2 exposed to 5, 10, and 15 minutes of UV, ∼0.6 for LEV-2 cultures irradiated for 1 hour or 8 hours, ∼0.4 for LEV-2 exposed to 2 hours or 4 hours of UV, and ∼0.8 for yeast cultures irradiated for 24 hours and for non-irradiated yeast cultures grown for 24 hours.

**Figure 4:**
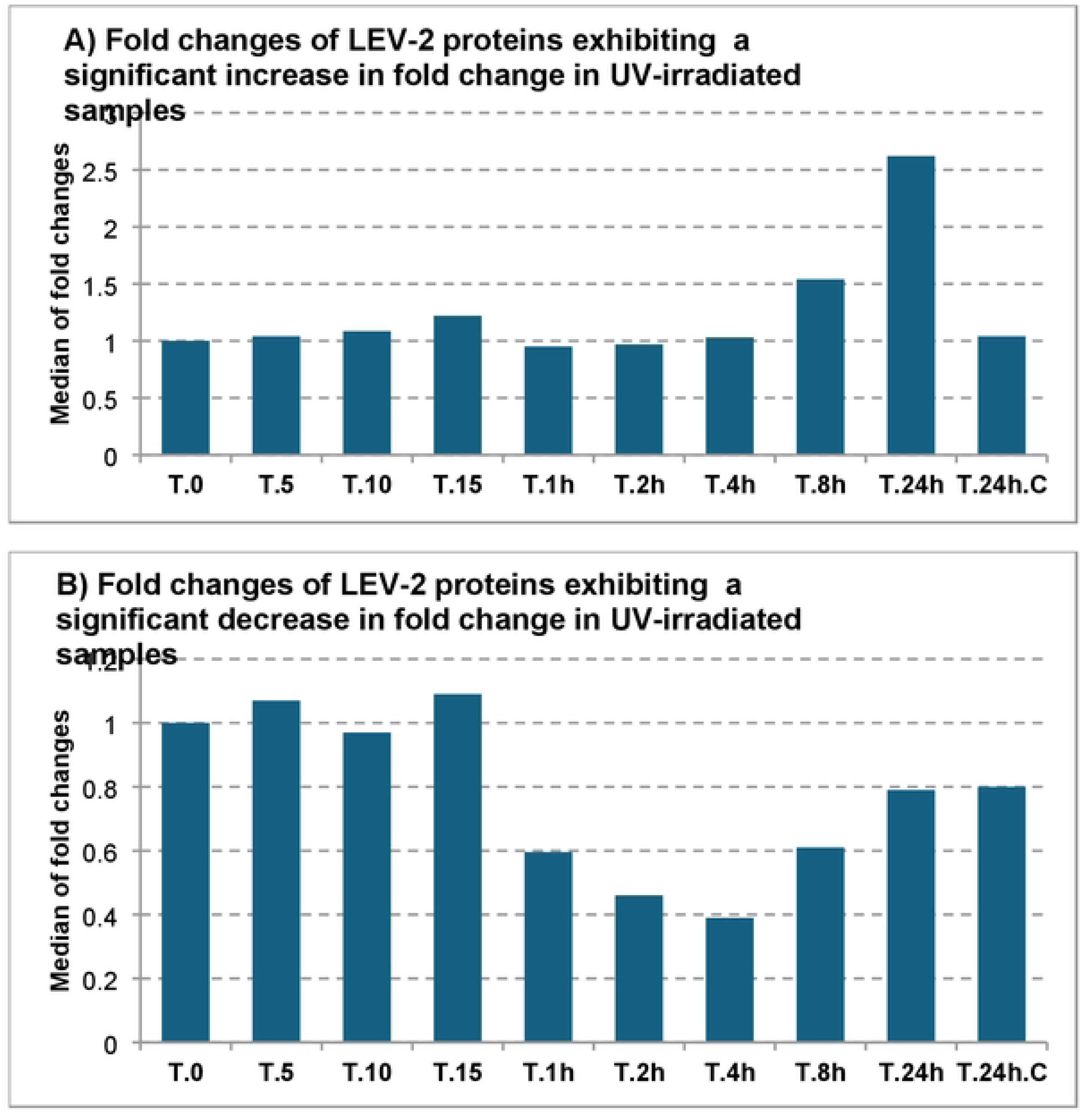
Expression profiles of proteins exhibiting a significant fold change. The figure (A) displays the median values of fold changes for 227 yeast LEV-2 proteins that exhibited 2-fold or higher increase in expression in at least one UV­ irradiated sample, relative to the non-irradiated control. The figure (B) shows the median values of expression fold changes for 279 yeast LEV-2 proteins that exhibited significant (2-fold) or higher decrease in fold change in at least one UV­ irradiated sample.

### Functional annotation of yeast LEV-2 proteome

The 752 yeast LEV-2 proteins quantified by MudPIT analysis were annotated for predicted biological function using the InterProScan [37] tool to assign Gene Ontology (GO) terms [31] to the proteins. In addition, KEGG tools [32] were used to assign predictions of biochemical pathways (KEGG pathways) and biological functions (KEGG Modules) to the quantified proteins. Statistical analysis based on Fisher’s exact test [39] was performed to identify GO terms, KEGG modules, and KEGG pathways over-represented among proteins showing fold change increases and fold change reductions in LEV-2 cultures exposed to UV-B radiation. For the dataset of 227 proteins exhibiting fold change increases in LEV-2 isolates exposed to UV-B, the over-represented GO terms (Table 1A) included GOs related to cellular transport systems, ribosome biogenesis, stress response, cellular signalling, and cellular respiration. Annotations based on KEGG pathways (Table 2A) and KEGG modules (Table 3A) also indicated that functions and pathways related to stress response, cellular signalling, and cellular respiration are over-represented in this dataset. In addition, the over-represented KEGG pathways also included pathways involved in metabolism of arginine, histidine, and mannose. Among the 279 LEV-2 proteins that exhibited fold change reductions in UV-exposed yeast cultures, the GO terms over-represented among these proteins (Table 1B) included GO terms related to protein biosynthesis, protein folding and degradation, ATP binding and synthesis, pentose-phosphate pathway, biosynthesis of nucleotides, and metabolism of certain amino acids such as glycine and serine. Annotation using KEGG pathways (Table 2B) and KEGG modules (Table 3B) identified that pathways involved in carbohydrate metabolism and ribonucleotide biosynthesis are over-represented in this dataset.

**Table 1:**
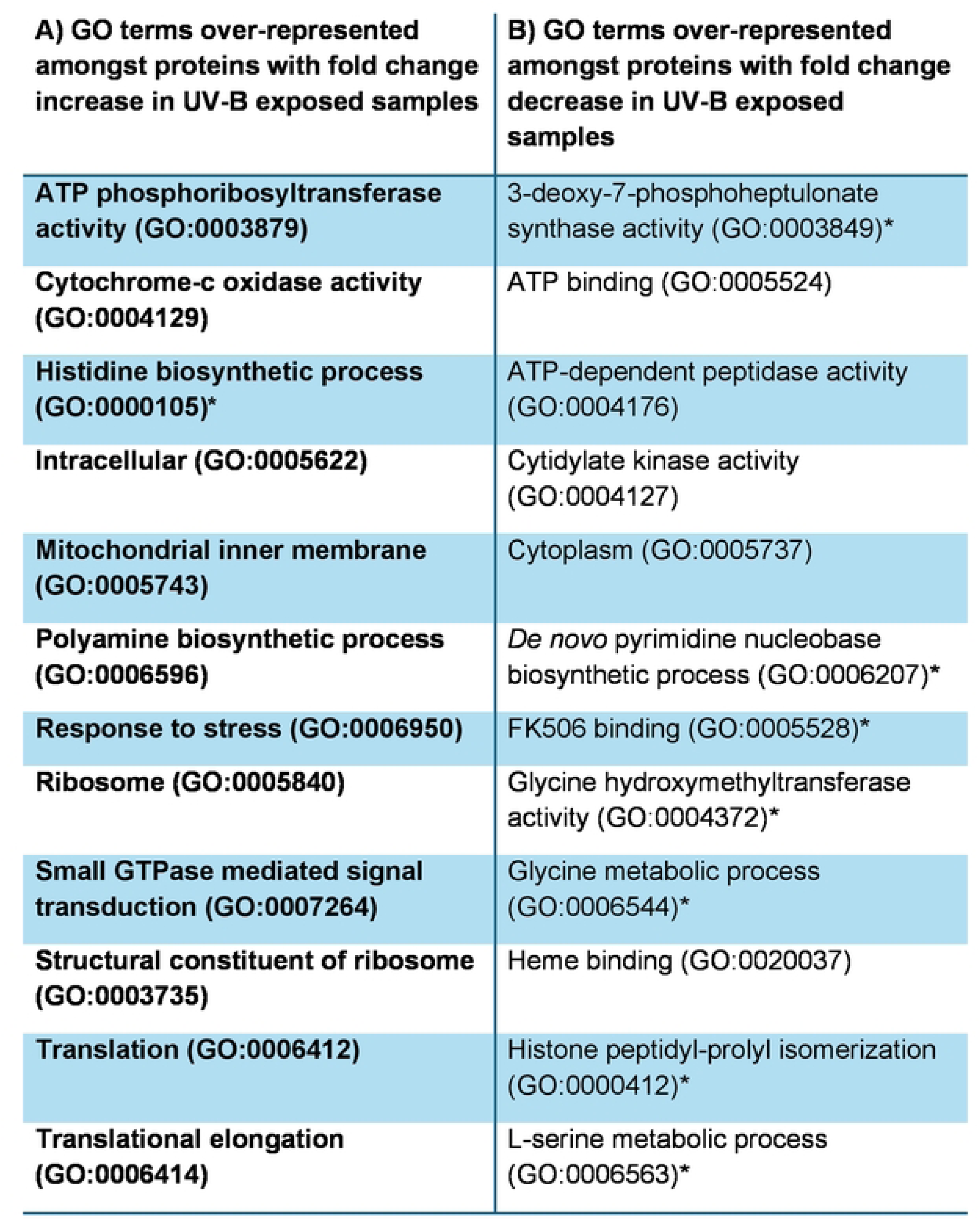
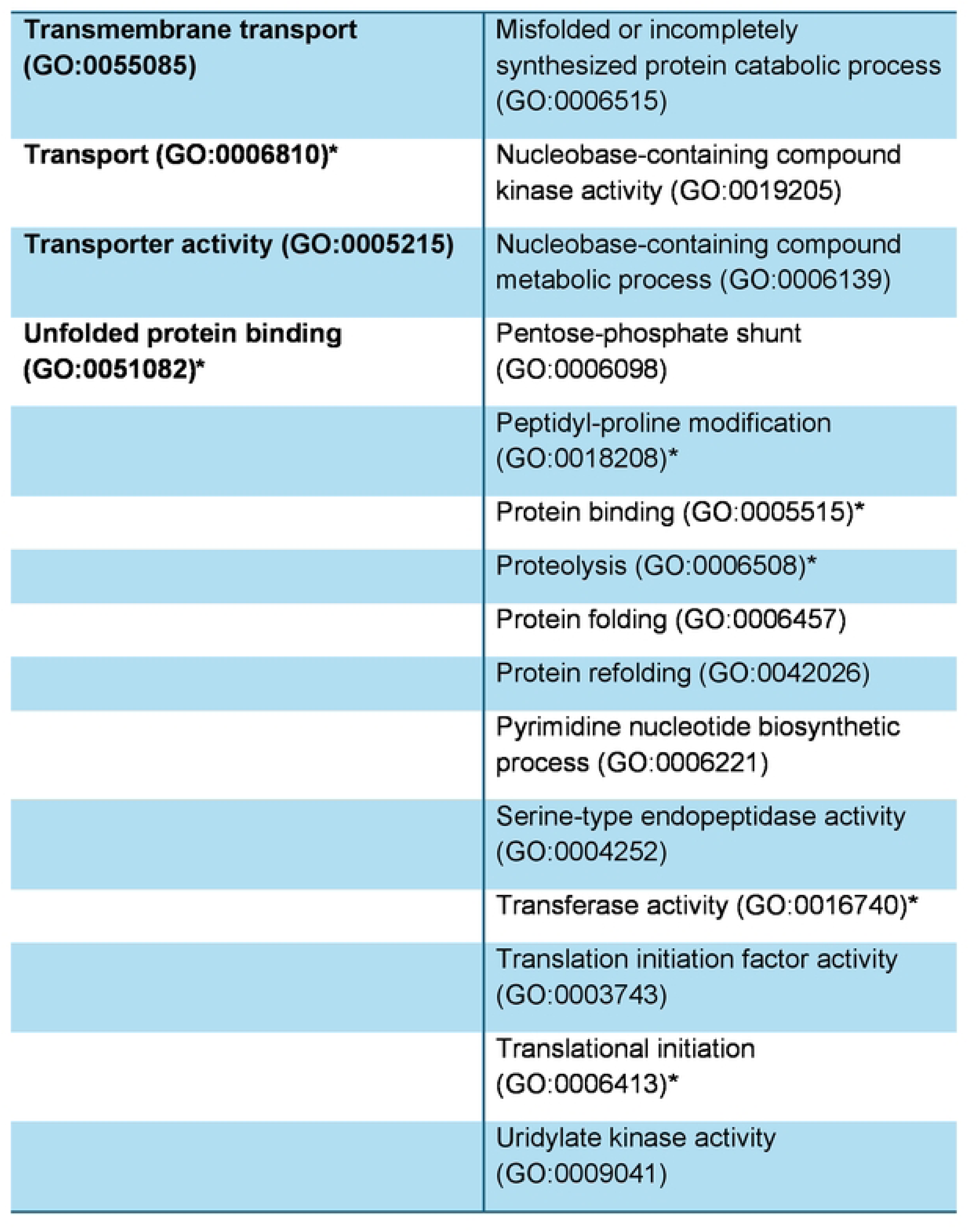
GO terms over-represented in datasets of LEV-2 proteins showing a significant fold in UV-B exposed yeast cultures. Column A lists GO terms over­ represented in a dataset of proteins exhibiting a significant fold change increase in UV-8 exposed yeast LEV-2 samples, while Column B lists GO terms over­ represented amongst proteins showing a significant fold change reduction. Terms marked by a star(*) were over-represented in a dataset of proteins with expression fold change of 4.0. The GO terms were considered over-represented if Fisher’s exact test resulted in p-value below 0.05, when frequencies of terms were compared between all proteins and proteins with fold-changes 2.0 or higher.

| **A) GO terms over-represented amongst proteins with fold change increase in UV-B exposed samples** | **B) GO terms over-represented amongst proteins with fold change decrease in UV-B exposed samples** |
| --- | --- |
| **ATP phosphoribosyltransferase activity (GO:0003879)** | 3-deoxy-7-phosphoheptulonate synthase activity (GO:0003849)* |
| **Cytochrome-c oxidase activity (GO:0004129)** | ATP binding (GO:0005524) |
| **Histidine biosynthetic process (GO:0000105)\*** | ATP-dependent peptidase activity (GO:0004176) |
| **Intracellular (GO:0005622)** | Cytidylate kinase activity (GO:0004127) |
| **Mitochondrial inner membrane (GO:0005743)** | Cytoplasm (GO:0005737) |
| **Polyamine biosynthetic process (GO:0006596)** | *De novo* pyrimidine nucleobase biosynthetic process (GO:0006207)* |
| **Response to stress (GO:0006950)** | FK506 binding (GO:0005528)* |
| **Ribosome (GO:0005840)** | Glycine hydroxymethyltransferase activity (GO:0004372)* |
| **Small GTPase mediated signal transduction (GO:0007264)** | Glycine metabolic process (GO:0006544)* |
| **Structural constituent of ribosome (GO:0003735)** | Heme binding (GO:0020037) |
| **Translation (GO:0006412)** | Histone peptidyl-prolyl isomerization (GO:0000412)* |
| **Translational elongation (GO:0006414)** | L-serine metabolic process (GO:0006563)* |
| **Transmembrane transport**<br>**(GO:0055085)** | Misfolded or incompletely synthesized protein catabolic process (GO:0006515) |
| **Transport (GO:0006810)\*** | Nucleobase-containing compound kinase activity (GO:0019205) |
| **Transporter activity (GO:0005215)** | Nucleobase-containing compound metabolic process (GO:0006139) |
| **Unfolded protein binding**<br>**(GO:0051082)\*** | Pentose-phosphate shunt (GO:0006098) |
|  | Peptidyl-proline modification (GO:0018208)* |
|  | Protein binding (GO:0005515)* |
|  | Proteolysis (GO:0006508)* |
|  | Protein folding (GO:0006457) |
|  | Protein refolding (GO:0042026) |
|  | Pyrimidine nucleotide biosynthetic process (GO:0006221) |
|  | Serine-type endopeptidase activity (GO:0004252) |
|  | Transferase activity (GO:0016740)* |
|  | Translation initiation factor activity (GO:0003743) |
|  | Translational initiation (GO:0006413)* |
|  | Uridylate kinase activity (GO:0009041) |

**Table 2:** KEGG pathways over-represented amongst the LEV-2 proteins showing a significant fold change in yeast LEV-2 exposed to UV-8. Column A lists KEGG pathways over-represented in a dataset of proteins exhibiting a significant fold change increase in UV-8 exposed yeast LEV-2 cultures, while Column 8 lists pathways over-represented amongst proteins showing a significant fold change decrease. The pathways were assigned by BlastKOALA search followed by KEGG pathway analysis. KEGG pathways marked by a star(*) were over­ represented in a dataset of proteins with expression fold change of 4.0 or higher. The KEGG pathways were considered over-represented if Fisher’s exact test resulted in p-value below 0.05, when frequencies of KEGG pathway terms were compared between all proteins and proteins with fold-changes 2.0 or higher.

| A) KEGG pathways over-represented amongst the proteins with a fold change increase in UV-B exposed LEV-2 | B) KEGG pathways over-represented amongst the proteins with a fold change decrease in UV-B exposed LEV-2 |
| --- | --- |
| Arginine and proline metabolism* | Cyanoamino acid metabolism* |
| Calcium signaling pathway* | HIF-1 signaling pathway |
| cAMP signaling pathway | One carbon pool by folate* |
| cGMP-PKG signaling pathway | Pyrimidine metabolism* |
| Fructose and mannose metabolism |  |
| Glutathione metabolism* |  |
| Histidine metabolism |  |
| Pentose and glucuronate interconversions |  |
| PI3K-Akt signaling pathway* |  |
| Ras signaling pathway |  |
| Ribosome |  |

**Table 3:**
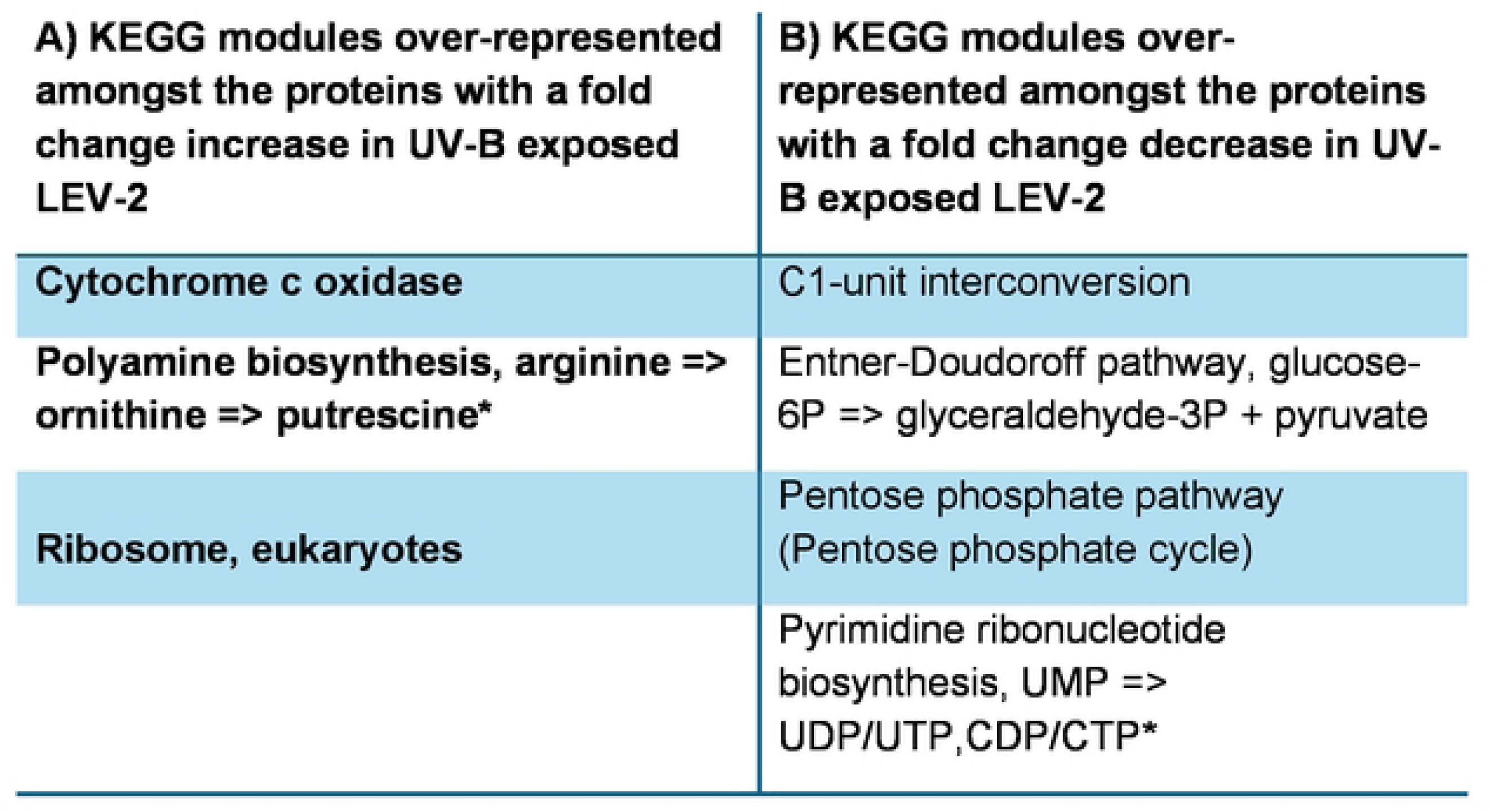
KEGG modules over-represented amongst the LEV-2 proteins showing a significant fold change in yeast LEV-2 exposed to UV-8. Column A lists KEGG modules over-represented in a dataset of proteins exhibiting a significant fold change increase in UV-8 exposed yeast LEV-2 samples. Column B shows KEGG modules over-represented in a dataset of proteins showing a significant fold change decrease. The KEGG modules were assigned by BlastKOALA search followed by KEGG module analysis. KEGG modules marked by a star(*) were over­ represented in a dataset of proteins with expression fold change of 4.0 or higher. The KEGG modules were considered over-represented if Fisher’s exact test resulted in p-value below 0.05, when frequencies of KEGG module terms were compared between all proteins and proteins with fold-changes 2.0 or higher.

### Yeast LEV-2 proteins involved in response to UV-B induced stress

Functional annotation of 751 quantified yeast LEV-2 proteins identified 105 proteins involved in cellular stress responses. Of these, four proteins were annotated as basic leucine zipper proteins, 22 as cellular signalling proteins, 17 as enzymes involved in biosynthesis of small molecule antioxidants, 22 as enzymatic antioxidants, six as enzymes involved in DNA repair, 24 as heat shock proteins. Four basic leucine zipper (bZip) containing proteins, similar to human Nrf2 and AP-1 proteins, were identified and quantified by MudPIT analysis using the database of fungal bZip proteins (Figure 5). The protein similar to XP_007274754.1 (bZip transcription factor of the fungus *C. gloeosporioides*) exhibited a significant fold change increase in LEV-2 yeast cultures exposed to 5 minutes, 1 hour, 2 hours, and 4 hours of UV-B. Proteins similar to GAP83664.1 (bZip protein from R. necatrix) and XP_748177.1 (bZip protein from A. fumigatus) showed significant fold change decreases in yeast cultures exposed to 4 hours and 8 hours of UV-B, while the bZip protein KFA50940.1 showed no significant fold changes in UV-B exposed cultures.

**Figure 5:**
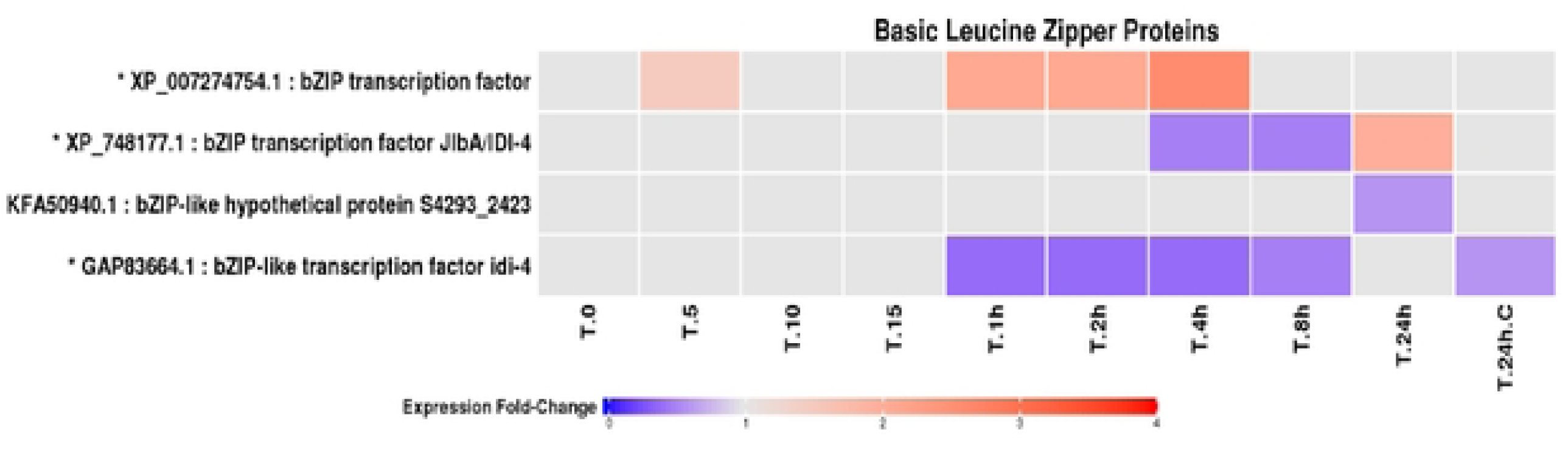
Fold change profiles of yeast LEV-2 bZip proteins: A heat-map of yeast isolate LEV-2 bZip protein fold changes in cultures exposed to UV-B for 5 minutes (T.5) to 24 hours (T.24h) and non-irradiated controls (T.O and T.24h.C). The proteins with fold change reduction in the UV-exposed samples are coloured blue, while proteins with fold change increases are coloured red. Proteins showing a fold change lower than 1.5 are coloured light-grey. The protein identifiers of the proteins showing 2-fold or higher fold change in at least one UV-exposed yeast culture are also marked with a star (*).

### Signalling proteins

Quantitative MudPIT analysis identified 22 signalling proteins (Figure 6). These were annotated as proteins involved in various signalling pathways including FoxO signalling [43], MAPK signalling [44], and RAS signalling [22]. Three 14-3-3 proteins were quantified, and all three exhibited fold change reductions in LEV-2 cultures irradiated for 1 hour to 4 hours. Four MAPK-signalling kinases were quantified, and all four exhibited fold-change increases in yeasts irradiated for 8 hours and 24 hours, while two MAPK-signalling kinases (similar to M6XZ23 and Spoli1_184897 proteins) also showed moderate (∼1.5-fold or lower) reductions in expression in LEV-2 cultures irradiated for 1 hour to 4 hours. Annotation identified two proteins involved in FoxO signalling, both of which exhibited moderate fold change increases in LEV-2 exposed to 8 hours and 24 hours of UV-B. Two Ras-related proteins were quantified; both of these proteins showed fold change increases in LEV-2 exposed to 24 hours of UV-B, while the Ras-related protein M7WXY7 also showed a moderate, ∼1.5-fold change increase in yeast cultures exposed to 15 minutes of UV-B. Four cell division control (Cdc) proteins were quantified; of these, two proteins belonging to the Cdc42 family showed fold change increases in yeasts exposed to 8 hours and 24 hours of UV-B, while two Cdc48 proteins exhibited fold change reductions in LEV-2 irradiated for 2 hours and 4 hours. A single Hippo-signalling protein was identified in this study; this protein did not show significant fold changes in UV-B irradiated LEV-2 cultures or in the controls. One calcium signalling protein, calmodulin, was identified and quantified and showed fold change increases in samples exposed to 1 hour to 4 hours of UV-B. The MudPIT analysis also identified three adenylate kinases and two other kinases for which detailed annotation could not be determined. The adenylate kinases showed fold change decreases in LEV-2 exposed to 1 hour to 24 hours of UV-B, the kinase Rhomi1_185026 exhibited fold change increases in yeast irradiated for 24 hours, and the kinase Rhomi1_141225 showed moderate fold change increases in LEV-2 irradiated for 5 to 15 minutes, 8 hours, and 24 hours.

**Figure 6:**
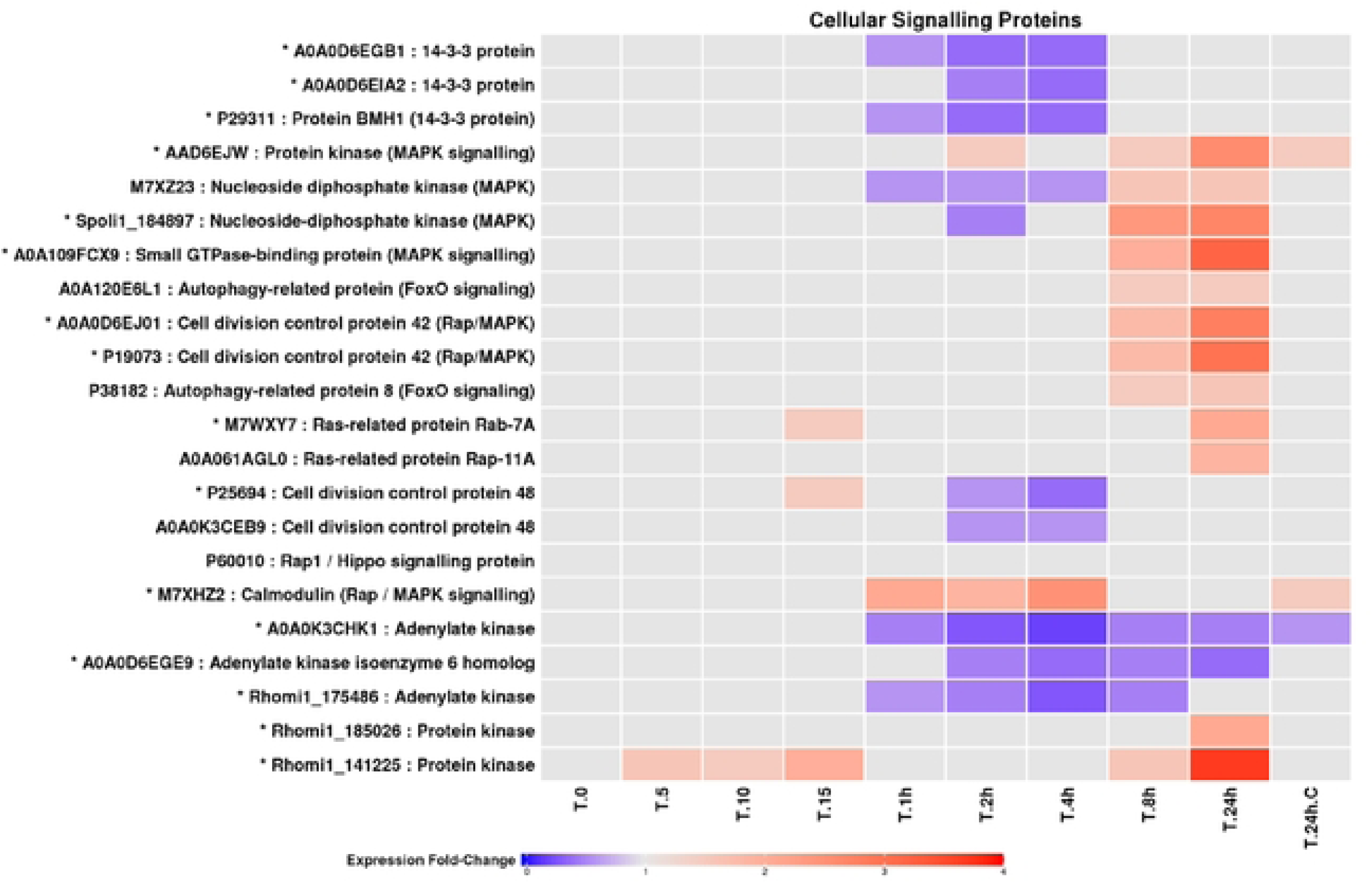
Fold change profiles of LEV-2 proteins involved in cellular signalling. A heat-map of fold changes of LEV-2 proteins involved in cellular signalling, cell cycle control and apoptosis. The fold changes are shown for LEV-2 cultures exposed to UV for 5 minutes (T.5) to 24 hours (T.24h) and in the cultures of non-irradiated controls (T.O and T.24h.C). The proteins showing a fold change reduction in the UV­ exposed samples are coloured blue, while proteins with fold change increase are coloured red. Proteins showing a fold change lower than 1.5 are coloured light-grey, and identifiers of proteins showing 2-fold or higher fold change in at least one UV­ exposed yeast culture are also marked with a star (*).

### Proteins involved in biosynthesis of antioxidants

MudPIT analysis of yeast isolate LEV-2 exposed to UV-B irradiation identified and quantified 17 proteins involved in biosynthesis of small-molecule antioxidants such as glutathione and ubiquinol (Figure 7). Hydroxyacylglutathione hydrolase (Rhomi1_37826), a protein involved in biosynthesis of glutathione [45], showed fold change reductions in LEV-2 cultures exposed to 4 hours to 24 hours of UV-B. Four glutathione transferase (GST) enzymes, which mediate phase II detoxification by catalysing the conjugation of xenobiotic functional groups with glutathione [45], were identified. The GST protein M7WWF5 exhibited fold change reductions in samples irradiated for 1 hour to 24 hours, while GSTs Rhomi1_153124 and Rhomi1_19319 showed moderate fold change decreases in yeasts irradiated for 2 hours and 4 hours. The GST protein Rhomi1_153124 also showed fold change increases in LEV-2 exposed to 15 minutes and 24 hours of UV-B. PdxS/SNZ family lyases and pyridoxine 4-dehydrogenases are involved in yeast biosynthesis of vitamin B6 and are associated with resistance to oxidative stress [46]. Four lyases of the PdxS/SNZ protein family were quantified by MudPIT analysis; these proteins showed fold change decreases in UV-irradiated samples (1 hour to 8 hours of UV-B). A single pyridoxine 4-dehydrogenase (Rhomi1_173574) was identified and quantified, and this protein showed no change in expression levels in UV-irradiated yeast cultures. Eight succinate dehydrogenase enzymes, involved in reduction of oxidized coenzyme Q10 to its reduced form (ubiquinol) [47], were identified and quantified in this study. Of these, protein P47052 showed no significant fold changes in any of the studied samples; A0A109FAY5 and Sporo1_19407 exhibited moderate, ∼1.5-fold change decreases in yeast LEV-2 exposed to 4 hours of UV-B; Spoli1_173565 exhibited moderate fold change decreases in samples exposed for 1 hour, 4 hours, and 8 hours; Rhomi1_182330 showed moderate fold change decreases in yeast irradiated for 2 hours and moderate fold change increases in samples exposed to 24 hours; and Sporo1_9115 and Rhomi1_167063 exhibited fold change increases in yeast cultures irradiated for 24 hours.

**Figure 7:**
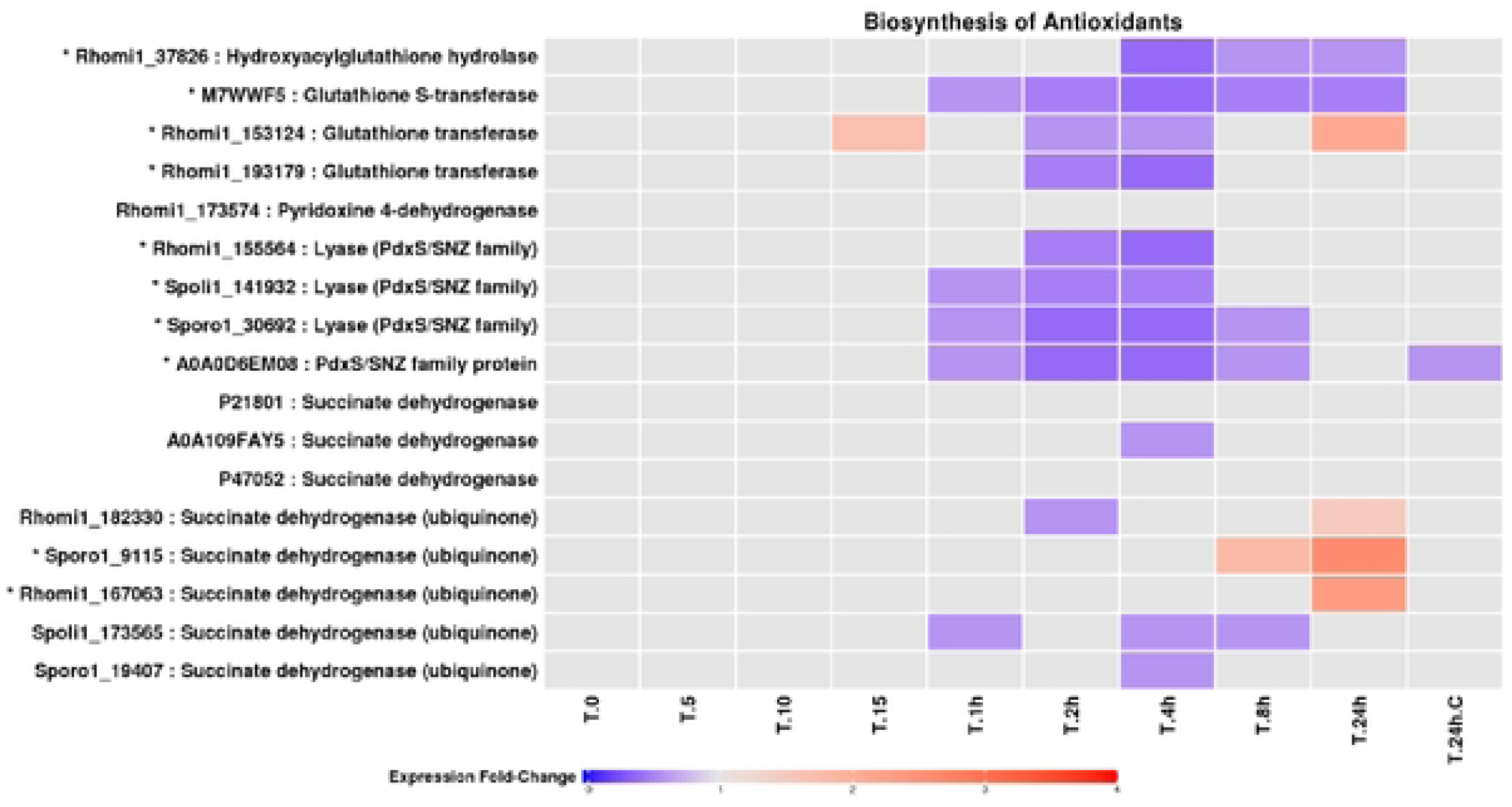
Expression profiles of LEV-2 enzymes involved in biosynthesis of antioxidants. **A** heat-map of the fold changes of yeast LEV-2 enzymes involved in the metabolism of small-molecule antioxidants. The LEV-2 cultures were exposed to UV for 5 minutes (T.5) to 24 hours (T.24h), and compared to non-irradiated controls (T.O and T.24h.C). The proteins showing a fold change reduction in the UV-exposed samples are coloured blue, while proteins showing a fold change increase are coloured red. The proteins exhibiting fold-changes lower than 1.5 are coloured light­ grey, and the proteins showing 2-fold or higher fold change in at least one UV­ exposed sample are marked with a star (*).

### Enzymatic antioxidants

The MudPIT study of UV-exposed yeast LEV-2 identified 22 enzymatic antioxidants, of which 6 showed significant fold change increases and 9 exhibited significant fold change decreases in yeast LEV-2 cultures exposed to UV-B (Figure 8). The identified antioxidants included superoxide dismutase (SOD), catalase, aldehyde dehydrogenase (ALDH), glutathione peroxidase (GTRx), isocitrate dehydrogenase (IDH), and cytochrome-c peroxidase (CCP) proteins, all of which are major enzymatic antioxidants in yeasts as well as animals [48]–[53]. One ALDH and one benzaldehyde dehydrogenase enzyme were identified and quantified in this study, and both enzymes showed significant fold change increases in LEV-2 isolates exposed to 24 hours of UV-B; the ALDH also exhibited moderate, ∼1.5-fold change increases in yeast exposed to 8 hours of UV-B. A single catalase was quantified; this enzyme showed moderate reductions in expression levels (∼1.5-fold change decrease) in LEV-2 cultures exposed to 2 hours to 8 hours of UV-B. Five CCP enzymes were quantified in yeast samples; two of these (Sporo1_16456 and A0A0D6ERS5) showed fold change reductions in LEV-2 cultures exposed to 1 hour to 24 hours of UV-B; CCP Sporo1_190216 exhibited fold change increases in samples exposed to 15 minutes, 8 hours, and 24 hours of UV-B and fold change decreases in samples exposed to 1 hour and 2 hours of UV; CCP Rhomi1_42978 showed fold change decreases in LEV-2 cultures exposed to 1 hour to 4 hours of UV and moderate fold change increases in cultures irradiated for 24 hours; and CCP Rhomi1_149378 showed fold change decreases in samples irradiated for 2 hours to 8 hours and increases in expression in LEV-2 cultures exposed to 15 minutes of UV-B. Two GTRx enzymes were identified by MudPIT proteomics analysis; the GTRx enzyme P38143 exhibited fold change reductions in yeasts exposed to 1 hour and 2 hours of UV, while the other GTRx (Rhomi1_146413) showed no changes in expression levels in UV-exposed yeast samples. The proteomics analysis identified two NAD⁺-dependent IDHs, both of which exhibited increases in fold changes in samples subjected to high doses of UV (8 to 24 hours of UV), and four NADP⁺-associated IDHs which displayed various degrees of fold change reductions in LEV-2 isolates exposed to 1 hour or longer UV irradiation. Four SOD enzymes were identified and quantified and showed significant fold change increases in LEV-2 samples irradiated for 24 hours. SODs Rhomi1_168331 and Spoli1_20960 also exhibited fold-change increases in yeast exposed to 8 hours, and SOD Rhomi1_86056 showed increases in fold changes in LEV-2 irradiated for 5 to 15 minutes and for 1 hour.

**Figure 8:**
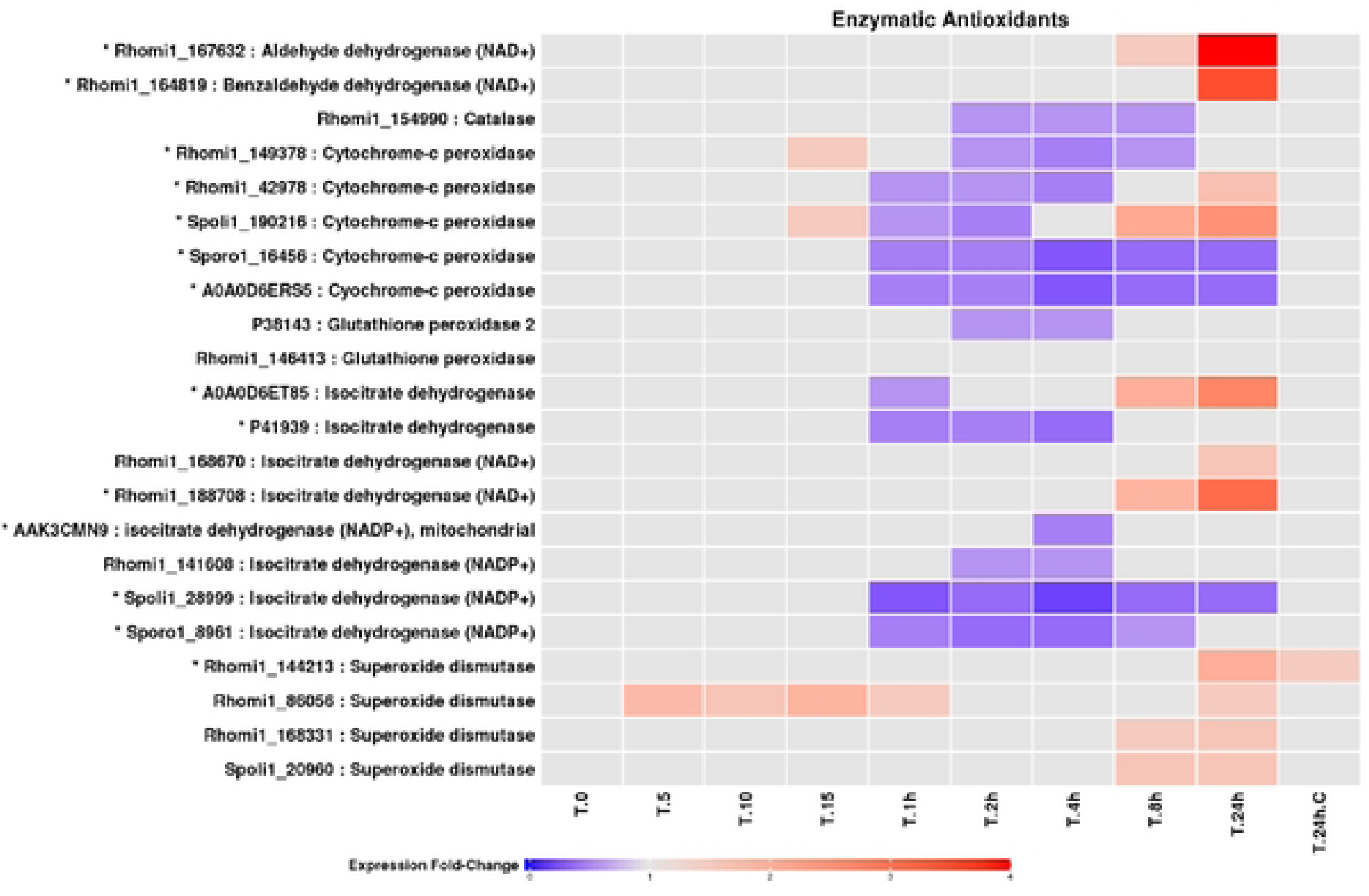
Fold change profiles of LEV-2 enzymatic antioxidants. The heat-map shows fold changes of enzymatic antioxidants of yeast LEV-2. The yeast cultures were exposed to UV for 5 minutes (T.5) to 24 hours (T.24h), and non-irradiated cultures were used as controls (T.O and T.24h.C). The proteins showing fold change reduction in the UV-exposed samples are coloured blue, while proteins with fold change increase are coloured red. The proteins exhibiting fold change lower than 1.5 are coloured light-grey, and identifiers of proteins showing significant (2-fold or higher) fold change in at least one UV-exposed sample are also marked with a star (*).

### Enzymes involved in repair and replication of DNA

Functional annotation of yeast LEV-2 proteins quantified by the MudPIT approach identified 6 enzymes involved in repair and replication of DNA (Figure 9). DNA ligase (Rhomi1_151258) and DNA-directed DNA polymerase (Rhomi1_90583) showed fold change increases in yeast samples irradiated for 24 hours. DNA ligase (Rhomi1_151268), DNA-directed DNA polymerase (Rhomi1_155605), and DNA helicase (A0A125PJD0) also exhibited moderate fold change reductions (fold change ∼1.5) in yeast samples exposed to 2 hours and 4 hours of UV.

**Figure 9:**
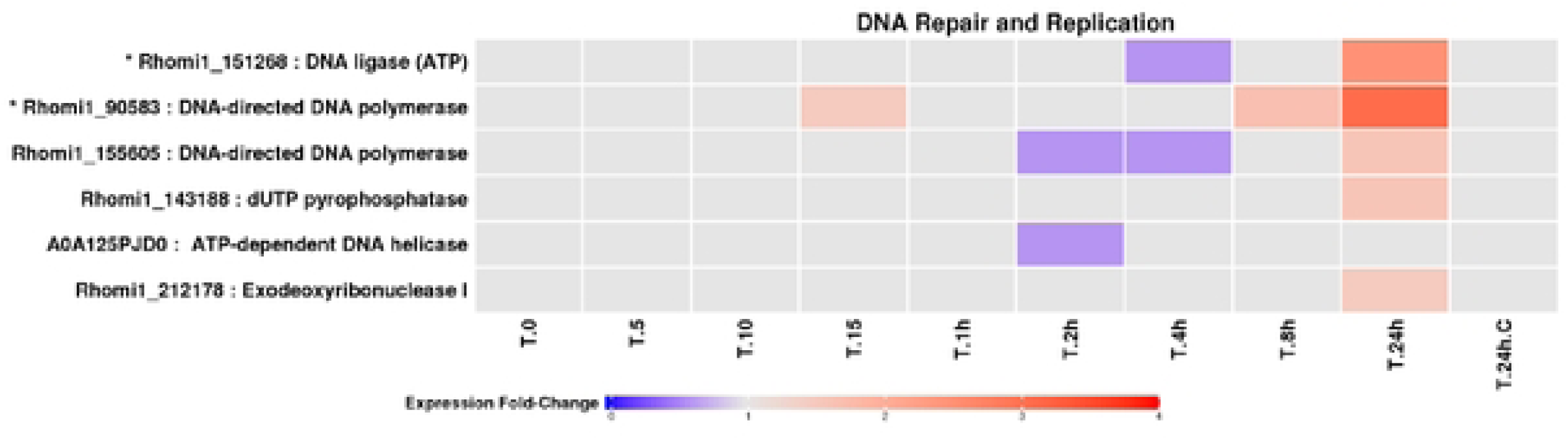
Fold change profiles of LEV-2 DNA repair and replication enzymes. **A** heat-map of fold changes of yeast LEV-2 enzymes involved in the repair and replication of DNA. The cultures of yeast LEV-2 were exposed to UV for 5 minutes (T.5) to 24 hours (T.24h), and compared to non-irradiated controls (T.O and T.24h.C). The proteins showing a fold change reduction in the UV-exposed samples are coloured blue, while proteins showing a fold change increase are coloured red. The proteins exhibiting fold-changes lower than 1.5 are coloured light-grey, and the identifiers of proteins showing significant, 2-fold or higher, fold change in at least one UV-exposed sample are marked with a star(*).

### Yeast heat-shock proteins and chaperonins

A total of 24 heat shock proteins (HSPs) were identified and quantified in this study (Figure 10). Two small, 10 kDa HSPs exhibited fold change increases in LEV-2 samples exposed to 8 hours and 24 hours of UV-B. The fold changes of all three identified chaperonin ATPases were increased in LEV-2 cultures exposed to 24 hours of UV-B. ATPase TCP-1 subunit theta (Rhomi1_150780) also exhibited fold change increases in LEV-2 irradiated for 4 hours and 8 hours, while ATPase TCP-1 subunits epsilon (Rhomi1_159882) and beta (A0A120E9R4) showed fold change reductions in LEV-2 irradiated for 2 hours. Six 60 kDa HSPs and nine 70 kDa HSPs were quantified; these proteins exhibited fold change reductions in LEV-2 cultures exposed to 1 hour to 8 hours of UV-B, and some of these proteins (Spoli1_199568, Rhomi1_3145, P19882, A0A0K3CKS4, A0A109FFK7, and P16474) also showed moderate fold change increases in yeasts irradiated for 24 hours. Of the four identified large, 90 kDa HSPs, all four exhibited high fold change increases in LEV-2 cultures irradiated for 24 hours, while three 90 kDa HSPs (A0A0D6EKA8, A0A109FF04, and Sporo1_21071) also displayed fold change increases in yeast irradiated for 8 hours, and HSP A0A109FKA3 also showed fold change reductions in samples irradiated for 2 hours and 4 hours.

**Figure 10:**
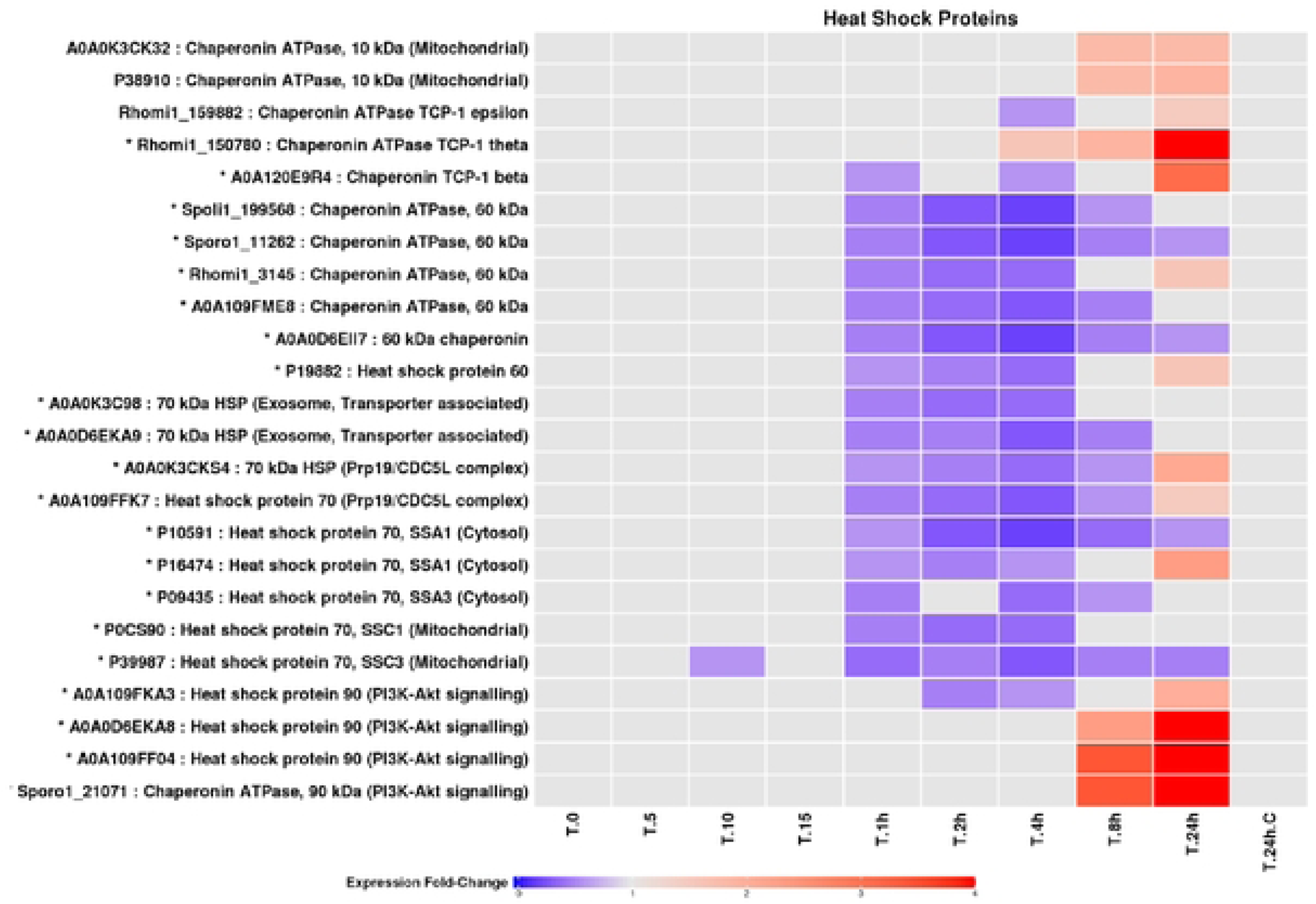
Fold changes of LEV-2 heat-shock proteins and chaperonins. A heat­ map of fold changes of yeast LEV-2 heat-shock proteins and chaperonins. The yeast LEV-2 cultures were exposed to UVR for 5 minutes (T.5) to 24 hours (T.24h), and compared to non-irradiated controls (T.Oand T.24h C). Proteins with fold change reduction in the UV-exposed cultures are coloured blue, while the proteins exhibiting a fold change increase are coloured red. Proteins with the fold changes lower than 1.5 are coloured light-grey, and the protein identifiers of proteins showing a significant, 2-fold or higher, fold change in UV-exposed samples are marked with a star(*).

## Discussion

The purpose of this study was to describe, at the proteome level, the stress response mechanisms of a UV-tolerant yeast, and to determine if bZip proteins play a role in this stress response. The *Sporobolomyces* yeast LEV-2, previously isolated from the leaves of Brazilian plants [30], was chosen for this study based on its high UV-tolerance (**Error! Reference source not found.**, **Error! Reference source not found.**), and because it was shown that cell extracts of LEV-2 cultures exhibited an increase in antioxidant, DPPH free radical quenching, activity during long-term exposure to UV-B (**Error! Reference source not found.**A). In addition, the previous analysis of this yeast placed it into Division Basidiomycota [30], unlike the commonly studied yeast *Saccharomyces cerevisiae,* which belongs to Division Ascomycota. The Ascomycota and Basidiomycota fungi are evolutionary distant, and are considered to have diverged ∼650 million years ago [54], [55]. Thus, this study enabled the comparison of stress response mechanisms between the major, evolutionary distant, divisions of fungi. When yeast LEV-2 was grown in liquid medium to mid-exponential phase of growth and exposed to UV-B irradiation for up to 24 hours. Solubilised proteins were extracted from the culture after 5 min, 10 min, 15 min, 1 hour, 2 hours, 4 hours, 8 hours and 24 hours of irradiation. A total of 751 proteins could be identified and quantified using TMT-labels and MudPIT high throughput proteomics, and the proteins were functionally annotated by InterProScan [37], KEGG tools [32], [33] and literature searches.

### UV-B irradiation induces the stress response of yeast LEV-2

In addition to causing direct DNA damage by inducing the formation of DNA photo-adducts [2], UV radiation stimulates the production of reactive, oxygen-derived, chemical species (ROS) in UV-exposed cells, and induces oxidative stress [3], [4]. The evidence of UV-caused oxidative damage has been found in multiple cell lines and *in-vivo* models. For example, UV-B radiation has been found to induce the production of ROS in human cell lines by enhancing the RS generating activity of catalase [4], to induce production of ROS in cyanobacteria, as detected by ROS-sensing probe 2’,7’-dichlorodihydrofluorescein diacetate [56], [57], and is considered also to be a major source of oxidative damage to yeast cells [58]. UV-A radiation was also found to cause single stranded DNA breaks associated with oxidative damage in mouse models [3] and in human skin [59]. In this study, a functional annotation of 227 proteins showing the fold change increase in yeast LEV-2 cultures exposed to UV-B (Figure 3A) identified a high number of proteins associated with stress-response and cellular signalling (Tables 1A to 3A). The increase in fold changes of these proteins in LEV-2 cultures exposed to 8 hours and 24 hours of UV-B (Table 3A) correlated with the increase of antioxidant activity of cell extracts of LEV-2 cultures exposed to 8 hours and 24 hours of UV-B, as measured by the DPPH free radical quenching assay (Figure 2A), suggesting that the yeast LEV-2 increased the rate of biosynthesis of antioxidant enzymes when exposed to UV-B.

While UV-stress responses of UV-tolerant *Sporobolomyces* yeasts have not been previously studied, the stress responses of yeast *S. cerevisiae* exposed to various environmental stresses, including heat-shock, pH changes, oxidants such as H_2_O_2_ and cadmium, hyper-osmotic shock, starvation, and ionizing radiation, have previously been quantified using microarrays [60]. These studies identified that the expression levels of large number of mRNA transcripts (∼900 genes) change when the yeast is exposed to stress, but the changes in transcript levels are transient, and often adjust to levels close to unstressed cells during the conditions of prolonged stress [60], [61]. The fold changes of proteins in UV-B exposed yeast LEV-2 cultures (Figure 4) match the patterns observed in these microarray studies of yeasts exposed to environmental stresses [60], [61], suggesting that the stress response mechanisms are conserved between yeast *S. cerevisiae* and *Sporobolomyces* yeast LEV-2. The conservation of stress response patterns between yeasts of Genus *Saccharomyces* (Division Ascomycota) and Genus *Sporobolomyces* (Division Basidiomycota) would indicate that the stress response mechanisms are conserved between fungal lineages that diverged approximately 650 million years ago [54], [55], and that the majority of Basidiomycota and Ascomycota fungi possess homologous stress response mechanisms. This is further supported by the recent proteomics study by Villegas et al (2014), which examined the stress response of yeast of the Genus *Rhodotorula* (Division Basidiomycota) exposed to oxidative stress induced by high levels of copper, and identified an increase in expression levels of stress response proteins such as heat shock proteins and superoxide dismutase in stressed yeasts [62]. This similarity of stress response in evolutionary distant groups of fungi is possibly because the stress signalling mechanisms evolved during the early evolution of eukaryotic life. Our previous phylogenetic studies of evolution of the Nrf2 pathway suggested that the bZip-protein based response to oxidative stress evolved in early eukaryotes as an adaptation to oxidative environment caused by the rising oxygen levels during the geological time [21]. The expression fold change patterns of stress response proteins, including the bZip proteins and other signalling proteins of UV-stressed *Sporobolomyces* yeast LEV-2 were further examined to evaluate if the stress response is conserved between the yeast *S. cerevisiae* and LEV-2, and the results are discussed in the following section.

### Stress response proteins of yeast LEV-2

MudPit analysis of UV-irradiated cultures of *Sporobolomyces* yeast LEV-2 quantified 105 proteins involved in cellular stress response. Stress response proteins were functionally annotated using the InterProScan, KEGG tools and manual annotation. In animals, the bZip transcription factor Nrf2 activates the transcription of a large number of genes [63], [64] encoding detoxification and antioxidant enzymes such as aldehyde dehydrogenases, glutaredoxins and thioredoxins, as well as enzymes involved in biosynthesis of glutathione [65]. The function of the bZip transcription factor Nrf2 was characterized by high-throughput technology such as Chip-seq and microarrays [63], [64], and its role in protection against oxidative stress was confirmed in cell lines and animal models [65]. For example, genetically modified mouse models deficient in Nrf2-encoding gene were found to be highly sensitive to carcinogens such as benzo[a]pyrene [66], to environmental pollutants such as diesel exhaust [10], and to toxicity of drugs such as acetaminophen [67]. In addition, pharmaceuticals that activate Nrf2-signalling, such as sulforaphane (SFN) and butylated hydroxyanisole, were shown to activate the transcription of Nrf2-regulated genes, such as *GST*, *GCLC* and *NQ01*, in wild type mouse models, but not in Nrf2-knockout models [68]. The activation of Nrf2 by SFN has also been shown to protect the human cell models and mouse animal models against UV-induced oxidative stress [12], [15], [69]. The bZip proteins homologous to vertebrate Nrf2 protein have also been studied in invertebrates such as the fly *Drosophila melanogaster* and the nematode *Caenorhabditis elegans*, and were found also to regulate the response to oxidative stress in these invertebrate models [70]–[72].

The genome of *S. cerevisiae* encodes several bZip transcription factors including Gcn4 and eight Yap proteins (Yap1 – Yap8). The bZip protein Gcn4 activates the biosynthesis of amino acids, and is involved in response to starvation, but also in UV-stress [23], as evinced from low UV-tolerance of Gcn4 knockout yeast models [22]. The bZip transcription factor Yap1 is a major regulator of oxidative stress response in *S. cerevisiae,* while Yap2, Yap5 and Yap8 proteins play a role in response to metal-induced stress, Yap4 and Yap6 in regulating response to osmotic stress, and the roles of Yap3 and Yap7 are currently unknown [26]. The function of Yap1 was inferred because *YAP1*-knockout yeasts, but not yeasts deficient in other *YAP* genes, were found to possess low activity of antioxidant enzymes such as superoxide dismutase and glutathione reductase, and display low adaptability to oxidative stress [73]. In addition, the Yap1-binding motif on DNA has been found in promotor regions of antioxidant genes such as *GSH1* and *TRX2* [26]. The homologs of Yap1 protein have been empirically validated in yeasts *Candida albicans* (Cap1 protein) and *Schizosaccharomyces pombe* (Pap1 protein) [74], suggesting that bZip-regulated response to oxidative stress is conserved between yeasts of Division Ascomycota. While the Yap1 homologs have not been experimentally confirmed in *Sporobolomyces* (Division Basidiomycota) yeasts, our previous bioinformatics study found that genomes of Basidiomycota fungi encode homologs of bZip proteins similar to animal bZip transcription factor Nrf2 [20], and the recent study by Jindrich and Degnan (2016), identified also that animal bZip-encoding genes evolved from the bZip-encoding gene of unicellular eukaryote ancestor of fungi and animals [75].

MudPIT proteomics analysis of *Sporobolomyces* yeast LEV-2 identified four bZip proteins similar to fungal bZip transcription factors (**Error! Reference source not found.**), suggesting that bZip proteins are conserved between *S. cerevisiae* and *Sporobolomyces* yeasts. The LEV-2 bZip protein, designated LEV-2_XP_748177.1, showed a significant fold change increase in UV-irradiated LEV-2 cultures (**Error! Reference source not found.**), suggesting that LEV-2_XP_748177.1 is a LEV-2 homolog of *S. cerevisiae* Yap1 and plays a role in stress signalling in the yeast LEV-2. This is because the expression of Yap1 is increased in *S. cerevisiae* cultures exposed to oxidants such as H_2_O_2_ and paraquat [76], while the expression of other *S. cerevisiae* Yap proteins is not changed significantly during oxidative stress [26], [77]. The fold change patterns of bZip protein LEV-2_XP_748177.1 in yeast LEV-2 exposed to long term UV-B irradiation (Figure 5) were followed by the fold change increase in stress-response proteins of yeast LEV-2 (**Error! Reference source not found.**), and these fold changes showed striking resemblance to changes in expression levels of bZip transcription factor Nrf2 in human liver cancer (HepG2) cells and rat renal epithelial cells exposed to electrophiles such as tBHQ and β-NF [78] or to heme [79]. In these studies, exposure to Nrf2 activator caused an increase in cellular concentration of Nrf2 after ∼30 minutes to 1 hour, followed by an increase in expression of Nrf2-regulated genes after 2 or more hours. While the observed protein fold changes in LEV-2 cultures exposed to UV-B suggest that the bZip protein LEV-2_XP_748177 of *Sporobolomyces* yeast LEV-2 is a homolog of *S. cerevisiae* bZip transcription factor Yap1 and of vertebrate bZip transcription factor Nrf2, other signalling proteins might also be involved in stress response in yeast LEV-2. For example, RAS/cAMP/PKA signalling and bZip protein Gcn4 are known to be involved in UV-response of yeast *S. cerevisiae* [22]. Thus, further studies, potentially involving gene deletion of LEV-2_XP_748177-encoding gene in yeast LEV-2 or gene-knockdown by siRNA, are required to exclude the possibility that the observed changes in proteome of yeast LEV-2 exposed to UV-B radiation are mediated by other signalling pathways. It should be noted also that the genome of yeast LEV-2 is currently not available, and the presence of bZip proteins in LEV-2 was inferred by database matching of tandem mass spectra generated from tryptic digests of LEV-2 proteins. Therefore, primary amino-acid sequence of discovered bZip proteins could not be established, and genome sequencing yeast LEV-2 is required to elucidate the DNA sequence of bZip protein encoding genes of yeast LEV-2. The assembly of LEV-2 genome would facilitate the phylogenetic analysis of LEV-2 bZip proteins and sequence similarity comparisons of LEV-2 bZip proteins to vertebrate and known yeast bZip proteins. In addition, the knowledge of DNA sequence of bZip-encoding genes of the yeast LEV-2 would allow the design of PCR primers for real-time PCR analysis of bZip-encoding mRNA transcripts in yeast LEV-2 exposed to oxidative stress, and the design of siRNAs for knock-down experiments. In addition to the XP_748177.1 like protein, three other bZip proteins were identified and quantified in the proteome of yeast LEV-2, and the expression levels of these proteins were reduced or unchanged in UV-B irradiated yeast cultures (**Error! Reference source not found.**). These proteins are likely to be homologs of yeast *S. cerevisiae* Yap proteins (Yap2 – Yap8) not involved in response to oxidative stress. This is because the yeast *S. cerevisiae* Yap2 – Yap8 proteins do not show significant fold changes in *S. cerevisiae* exposed to oxidants [26], [60], [61], [77], and do not regulate the response to oxidative stress [24]. For example, Yap5 protein is involved in iron metabolism [80], Yap6 regulates the response to osmotic stress, and functions of Yap3 and Yap7 are currently unknown [26].

### Signalling and apoptosis related proteins

The proteomics study of LEV-2 yeast identified 22 proteins associated with different cellular signalling pathways. These proteins were further classified based on the pathway:

#### The 14-3-3 proteins

The 14-3-3 proteins are involved in numerous cellular processes, including signal transduction, cell-cycle control and apoptosis, and signalling roles of these proteins are an active field of research [81]. Three 14-3-3 proteins were identified in this study, and the fold changes of these proteins were significantly reduced by moderately long UV-B irradiation (1 hour to 4 hours of UV-B exposure). This reduction in expression co-occurs with increase in cell death observed in samples under 1 hour to 4 hours of UV-B (**Error! Reference source not found.**), suggesting the link between levels of 14-3-3 proteins and cell death, possibly by UV-induced apoptosis. This is in agreement with studies of Zhang et al. (1999), who identified a strong correlation between expression of 14-3-3 proteins and apoptosis in HeLa cell line model [82]. It is currently unknown whether 14-3-3 proteins play identical roles in human cell line models and in yeasts, and different stresses were reported to affect the expression of 14-3-3 proteins in yeast *S. cerevisiae* in different fashion, depending on the source of stress. For example, Yoshimoto et al. (2002) found that calcium induced stress caused the reduction in expression levels of 14-3-3 proteins in *S. cerevisiae*, as measured by microarrays [83]. The similar microarrays-based study by Gasch et al. (2002) identified that changes in levels of 14-3-3 proteins depend on the source of stress, with protein levels increased in yeast *S. cerevisiae* exposed to heat-shock, and reduced in the sample exposed to H_2_O_2_ [61].

#### MAPK signalling proteins

The mitogen-activated protein kinase (MAPK) signalling pathway mediates transduction of extracellular signals, and is essential for multiple cellular functions, such as cell differentiation, proliferation, initiation of apoptosis and adaptation to environmental stresses [44]. The basic assembly of MAPK signalling comprises three kinases - MAPK kinase kinase (MAPKKK), MAPK kinase (MAPKK), and MAP kinase (MAPK) - that transduce extracellular signal by sequential phosphorylation reactions. MAPKKK phosphorylates MAPKK, which in turn phosphorylates MAPK, and the phosphorylated MAPK phosphorylates transcription factors such as Sko1 which control transcription of genes involved in stress response [84]. This three-component module is conserved between yeasts and animals [84], and the increase in expression of MAPK kinases has been associated with increase in resistance to oxidative stress in yeasts *S. cerevisiae* and *S. pombe* [74], invertebrates such as *C. elegans* [85], mouse animal models and human cell lines [84], [86]. For example, *S. cerevisiae* and *S. pombe* yeasts deficient in MAPK kinase encoding genes are hypersensitive to salt-induced stress, heat shock and nutritional limitation [74]; in *C. elegans,* oxidative stress induced by sodium arsenite, paraquat, or t-butyl peroxide increases the expression of MAPK kinase p38, while the deletion of gene encoding the MAPK kinase SEK-1 increases sensitivity to oxidative stress [85]; human HeLa cells exposed to H_2_O_2_-induced oxidative stress exhibit an increase in expression of multiple MAPK kinases, and the inhibition of extracellular MAPK signalling kinases increases the apoptosis rate in stressed cells [86].

The cell division control proteins 42 (Cdc42) were shown to be involved in activation of yeast high osmolarity glycerol (HOG) MAPK pathway [87]. The HOG pathway involves a series of kinases which activate Hog1 MAP kinase in response to severe osmotic stress, and the mutation of yeast *S. cerevisiae* gene encoding Cdc42 proteins has been shown to inhibit the yeast response to osmotic stress, while mutants over-expressing Cdc42 proteins have been found to exhibit an increase in stress response [87]. This is because Cdc42 proteins interact with, and activate, the Ste20 MAP4K component of HOG pathway in yeast, and this interaction is essential for signal transduction during osmotic stress [88]. This function of Cdc42 proteins in MAPK pathway is conserved between yeast and mammals, and it has been shown that stress-activated MAPK pathway kinases such as JNK1 and p38 are activated by the Cdc42 protein in human tissue extracts [89] as well as monkey kidney cell (COS-1) cultures [90].

Four kinases associated with MAPK signalling pathways, and two Cdc42 proteins, were identified by MudPIT analysis of *Sporobolomyces* yeast LEV-2. These six proteins showed fold change increases in yeast cultures exposed to long term, 8 hours or longer, UV-B radiation (**Error! Reference source not found.**). These results suggest that a MAPK signalling is activated by UV-B radiation in yeast LEV-2, and that these signalling pathways are evolutionary conserved between Ascomycota yeasts such as *S. cerevisiae* and *S. pombe*, and the *Sporobolomyces* yeast LEV-2. While MAPK pathway has not yet been studied in *Sporobolomyces* yeasts, this pathway was empirically characterized and found to mediate a response to stress in Basidiomycota fungi *Cryptococcus neoformans* exposed to fungicide fludioxonil and to high levels of NaCl [91]. In this study, the increase in MAPK signalling kinases was found to correlate with the fold change increase of enzymatic antioxidants, such as SODs (**Error! Reference source not found.**), and with the fold change increases of certain heat shock proteins (**Error! Reference source not found.**) observed in LEV-2 cultures exposed to 8 hours and 24 hours of UV-B radiation. In addition, the cell extracts of LEV-2 cultures irradiated for 8 hours and 24 hours also showed the increase in DPPH free radical scavenging activity (**Error! Reference source not found.**). These results, and the assumption that MAPK-mediated stress response is conserved across all eukaryotes [84] indicate that MAPK signalling is likely involved in activating stress response in UV-irradiated yeast LEV-2. It is however, known that other signalling pathways, such as Yap1 signalling [26], are also involved in yeast stress response, and further empirical studies, such as inactivation of genes encoding MAPK kinases in yeast LEV-2, are required to establish the importance of MAPK signalling in yeast LEV-2 in comparison to other signalling pathways, such as the Yap1 pathway.

#### FoxO signalling

FoxO signalling proteins represent a subfamily of the forkhead family of transcription factors, and are highly conserved across all animal phyla, with orthologs discovered in cnidarian *Hydra vulgaris,* worm *C. elegans,* fly *D. melanogaster*, mouse and rat models and human cell lines[92]. These proteins have been associated with resistance to oxidative stress, control of life-span, regulation of cell cycle arrest, and induction of apoptosis [92]. In the nematode *C. elegans,* FoxO signalling is mediated by the forkhead transcription factor Daf-16, and the increase in expression of this protein was shown to increase the worm life-span and resistance to oxidative stress [93]. The increase in FoxO signalling was found also to mediate stress response in a mammalian cell line: H_2_O_2_-induced oxidative stress caused the fold change increase of the p66shc protein, mammalian homolog of Daf-16, while cells deficient in p66shc-encoding gene were found to be highly sensitive to H_2_O_2_ [94].

The yeast *S. cerevisiae* forkhead proteins HCM1, FKH1 and FKH2 proteins are homologs of animal FoxO proteins, and have been shown to regulate stress response and longevity of *S. cerevisiae* [43], [95], [96]. For example, the study by Postnikoff et al. (2012) found that *S. cerevisiae* mutants deficient in genes encoding FKH1 and FKH2 proteins had a shorter life-span and lower resistance to H_2_O_2_ than wild-type yeasts, while *S. cerevisiae* genetically modified to overexpress FKH proteins showed an increase in life-span and resistance to H_2_O_2_ [43]. The study by Maoz et al. (2015) also found that *S. cerevisiae* genetically modified to overexpress FoxO homolog HCM1 had a high rate of transcription of genes encoding catalase and superoxide dismutase enzymes; in addition, these yeasts were highly resistant to oxidative stress induced by H_2_O_2_, and had a higher lifespan than wild-type yeasts [95].

Two FKH proteins were identified and quantified in this study (**Error! Reference source not found.**), and levels of these proteins were moderately increased (fold change ∼1.5) in the yeast LEV-2 cultures exposed to long-term UV-B radiation (8 hours and 24 hours). This increase in expression of FKH proteins co-occurred with the increase of free radical quenching activity of cell extracts of LEV-2 yeast cultures exposed to UV-B radiation (**Error! Reference source not found.**) and with a significant fold change increase of superoxide dismutase enzymes (**Error! Reference source not found.**). These expression patterns match the results of previous studies that quantified the expression of FKH proteins of yeast *S. cerevisiae* exposed to H_2_O_2_-induced stress [43], [95], and imply that a fold change increase in FKH proteins is associated with yeast LEV-2 resistance to UV-induced oxidative stress. These results also suggest that the FoxO pathway is conserved between yeast *S. cerevisiae* and *Sporobolomyces* yeasts such as LEV-2.

#### Ras signalling proteins

In yeasts, such as the model yeast *Saccharomyces* cerevisiae, the Ras signalling pathway controls the DNA damage-independent response to UV-induced stress. This stress response pathway is comprised of membrane associated Ras proteins that activate the adenylate cyclase enzymes to stimulate the production of cyclic AMP (cAMP). The increased cytosolic concentration of cAMP activates a protein kinase A (PKA) controlled phosphorylation cascade that increases the translation of bZip transcription factor Gcn4 [22], and leads to induction of genes involved in biosynthesis of amino acids [23]. The yeast Ras signalling pathway is considered to be homologous to mammalian UV-response pathway that includes Ras associated proteins Ha-Ras and Raf-1 as well as transcription factors such as NF-kB and AP-1 [22], [97], and is distinct from yeast response to DNA damage, which is mediated by the DNA damage responsive protein kinase Dun1 [28]. This is because Ras signalling involves membrane associated Ras proteins rather than DNA damage sensing kinases. In addition, Ras signalling is involved in regulation of amino acid biosynthesis, but not in regulation of DNA repair, and transcription of Ras associated proteins is induced by UV in *S. cerevisiae* strains deficient in Dun1-encoding gene [22]. While primarily associated with response to starvation and with induction of amino acid biosynthesis [98], Ras signalling was shown to also regulate the protection against UV-induced stress in the yeast *S. cerevisiae*. This is because the *S. cerevisiae* strains deficient in gene encoding transcription factor Gcn4 are highly sensitive to UV irradiation, while the strains engineered to constitutively over-express Gcn4 are highly resistant to UV [22].

In this study, the Ras-associated proteins Rap-7A and Rap-11A of *Sporobolomyces* yeast LEV-2 showed a moderate, 1.5-fold, increase in LEV-2 yeast cultures exposed to 24 hours of UV-B (**Error! Reference source not found.**), suggesting that Ras signalling is enhanced in yeast LEV-2 during long-term UV-B irradiation. This increase in fold change co-occurred with the low death rate observed for LEV-2 cultures exposed to 8 hours and 24 hours of UV-B (**Error! Reference source not found.**), suggesting that the Ras-associated proteins of yeast LEV-2 induce the adaptation to UV-B stress, and that the function of Ras-associated proteins is conserved between *S. cerevisiae* and the yeast LEV-2. The observed reduction in yeast LEV-2 cell death is, however, also associated with an increase in expression of enzymatic antioxidants (**Error! Reference source not found.**) and other signalling pathways involving bZip transcription factors (**Error! Reference source not found.**), FoxO signalling, and MAPK signalling (**Error! Reference source not found.**). Thus, further research is required to quantify the contributions of individual signalling pathways to the stress response of the UV-exposed yeast LEV-2.

The Ras-related calcium-binding protein calmodulin is an important component of stress signalling in the yeast *S. cerevisiae*, and yeast strains deficient in the calmodulin-encoding gene have been shown to be sensitive to stress induced by elevated levels of ions such as OH^−^, Mn^2+^, Na^+^ and Li^+^ [99]. The calmodulin and Ca^2+^/calmodulin-dependent phosphatase calcineurin mediate the response to elevated levels of cytosolic Ca^2+^ caused by oxidative stress, osmotic shock and other sources of stress [99], [100]. Calcineurin responds to stress-associated elevation of Ca^2+^ levels in the cell by dephosphorylating the transcription factor Crz1p. The dephosporilated Crz1p translocates to the nucleus, where it activates transcription of genes encoding numerous proteins involved in cell survival [99]. For example, the microarray study of Yoshimoto et al. (2002) identified that over 150 genes are under transcriptional control by Crz1p in the yeast *S. cerevisiae*, including genes involved in control of ion transport and homeostasis, cell wall synthesis/maintenance, lipid and sterol metabolism, vesicle transport, and cellular signalling [83]. The stress response function of calmodulin signalling is conserved between *S. cerevisiae* and the pathogenic yeast *Candida albicans*, and it was shown that *C. albicans* deficient in genes encoding calmodulin-dependent protein kinases is highly sensitive to H_2_O_2_-induced oxidative stress [101]. This pathway, however, is not involved in the stress response of fungus *Aspergillus nidulans* [100], indicating that the function of calmodulin is not conserved across the different fungal phyla. In this study, the LEV-2 calmodulin protein (**Error! Reference source not found.**) showed a fold change increase in yeast cultures exposed to 1 hour to 4 hours of UV-B, and showed the pattern similar to LEV-2 bZip transcription factor, which also showed a fold change increase in UV-B exposed LEV-2 cultures (**Error! Reference source not found.**). Studies of different cell models such as hepatocytes, lymphocytes and endothelial cells, showed that oxidative stress disrupts the function of cellular Ca^2+^ transporters and increases the concentration of Ca^2+^ in the cytosol [102]. Thus, the observed fold change increase in the yeast LEV-2 calmodulin protein suggests that the calmodulin activity is increased in the yeast LEV-2 in response to an increase in cytosolic concentration of Ca^2+^ due to UV-induced oxidative stress. This would indicate that calmodulin is an early sensor of cellular stress in yeast LEV-2, and the function of this protein is conserved between yeast *S. cerevisiae* and *Sporobolomyces* yeast LEV-2.

#### Cell cycle control and apoptosis

Yeasts, and other unicellular organisms, were traditionally considered not to possess the mechanisms for induction of programmed, apoptotic, cell death, but the expression of mammalian proapoptotic genes such as *BAX* and *TP53* was shown to induce the apoptotic cell death in yeast *Saccharomyces cerevisiae* [103], [104]. In addition, it was shown that a moderate concentration of H_2_O_2_ (3 – 5 mM) induces apoptotic cell death in *S. cerevisiae*, while a high concentration of H_2_O_2_ (180 mM) causes necrotic cell death [103]. The discovery of the *S. cerevisiae* caspase *YCA1* and the yeast apoptosis-inducing factor-1 (*AIF1*), and studies that demonstrated that cell death is reduced in *S. cerevisiae* yeasts deficient in genes encoding these proteins, further confirmed that certain apoptotic cell death mechanisms are conserved between yeasts and animals [105]. The yeast LEV-2 homologs of genes encoding Yca1 and Aif1 proteins were not discovered in this study, possibly because amino acid sequences of these proteins in yeast LEV-2 are significantly different from the proteins of fungi for which proteomes were available, and which were used for database matching of tandem mass spectra generated from the tryptic fragments of LEV-2 proteins. The cell division proteins 48 and Ras associated proteins were, however, successfully identified and quantified (**Error! Reference source not found.**), and these proteins are known to be involved in apoptosis in yeast *S. cerevisiae* [106], [107].

The cell division proteins 48, referred to as Cdc48 in yeasts, and p97 proteins in animals, are multipurpose proteins conserved across fungi and animals, and essential for growth of the yeast *S. cerevisae* [108]. These proteins have been associated with numerous functions including protein degradation, protein aggregation, control of cell cycle and apoptosis, and transcription and replication of DNA [108]. The yeast Cdc48 proteins, and animal homologs, were also shown to play a role in endoplasmic reticulum (ER) stress; these proteins are involved in extraction of misfolded proteins from the ER into the cytosol, where the misfolded proteins are subsequently degraded [109]. Mutation in the *S. cerevisiae CDC48* gene has been shown to increase the rate of apoptosis in yeast [107], and an increase in apoptosis was also shown in human cell line, and in the fly *D. melanogaster* when the translation of VCP protein, the animal homolog of Cdc48, is knocked-down by siRNAs [110]. The fold change reduction of Cdc48 proteins of yeast LEV-2 was observed in yeast cultures exposed to 2 hours and 4 hours of UVR (**Error! Reference source not found.**), coinciding with the fold change decrease in proteins associated with catabolism of misfolded proteins, protein folding and protein refolding (Table 1A, Table 3B). In addition, the cell death rate of yeast LEV-2 cultures exposed to 2 hours and 4 hours of UV-B was considerably higher than the death rate of LEV-2 cultures exposed to 8 hours or 24 hours of UV-B (the time period where the Cdc48 proteins do not show fold change decrease). These results suggest that Cdc48 proteins are linked to misfolded protein stress, and possibly to apoptosis, in *Sporobolomyces* yeast LEV-2, and imply that function of Cdc48 proteins is conserved between *Basidiomycota* fungi and *Ascomycota* fungi.

In addition to the UV response, Ras and Ras-related (Rap) proteins have been associated with control of cell cycle and apoptosis in yeast *S. cerevisiae* [111], and constitutive high expression of Ras signalling proteins was demonstrated to reduce the lifespan, and induce the apoptotic cell death, of yeast *S. cerevisiae* [112], [113]. The study by Gourlay and Ayscough (2006) found that constitutive activation of Ras signalling, induced by knock-out of genes that encode the Ras regulating proteins Sla1p or End3p, leads to an increase in cAMP levels in yeast, followed by the accumulation of RS in the cytosol and apoptotic cell death [113]. Similar results were reported by Heeren et al. (2004), who found that mutations in *RAS* genes of *S. cerevisiae* led to redox imbalance characterized by the excretion of cytosolic glutathione, elevated RS concentration, reduction in yeast life-span, and apoptotic cell death [112]. The activation of apoptosis by Ras/cAMP/PKA signalling was also demonstrated for the pathogenic yeast *Candida albicans*, where the mutations that block Ras signalling, such as deletions of *RAS1* and *CDC35* genes, were found to suppress the apoptotic response to acetic acid or H_2_O_2_, while mutations that stimulate Ras signalling, such as *RAS1^val13^* mutation, accelerated the rate of apoptosis when *C. albicans* was exposed to acetic acid or H_2_O_2_ [114] . Unlike yeasts, the constitutive expression of Ras signalling has been associated with carcinogenesis in animals. For example, mutations in *HRAS*, *KRAS* and *NRAS* oncogenes that lead to constitutive transcription of Ras-controlled genes have been associated with human cancers and carcinogenesis in animal models [115].

The MudPIT study of the UV-B irradiated *Sporobolomyces* yeast LEV-2 quantified six Ras-associated proteins (**Error! Reference source not found.**). The levels of these proteins, however, did not show significant fold change increase in LEV-2 exposed to 1 hour - 4 hours of UV-B irradiation, which is the time period of high cell death rate (**Error! Reference source not found.**). This result indicates that UV-B induced cell death of yeast LEV-2 is not due to over-expression of Ras signalling, and is likely caused by other mechanisms, such as UV-induced generation of RS leading to oxidative stress. In addition, the yeast LEV-2 death rate (**Error! Reference source not found.**) showed a marked decrease in yeast LEV-2 cultures exposed to 8 hours of UV-B and 24 hours of UV-B; this reduction in cell death coincided with the fold change increases in enzymatic antioxidants (**Error! Reference source not found.**), 10 kDa and 90 kDa heat shock proteins (**Error! Reference source not found.**), and proteins involved in the MAPK, Ras and FoxO signalling pathways (**Error! Reference source not found.**). Furthermore, the antioxidant activity of cell extracts of LEV-2 yeasts, as measured by DPPH free radical quenching was also highly elevated after 8 hours and 24 hours of UV-B exposure (**Error! Reference source not found.**). These results indicate that the increase in antioxidant activity is linked with reduction in yeast LEV-2 cell death, suggesting that the primary cause of cell death of yeast LEV-2 exposed to UV-B is oxidative stress, possibly leading to apoptosis, rather than direct, UV-inflicted, DNA damage. While this is consistent with previous studies that identified that moderate oxidative stress induces apoptosis rather than necrosis [103] in yeast *S. cerevisiae*, it should be noted that measurement of yeast viability after UV-B irradiation is not sufficient to differentiate between apoptotic cell death and necrotic cell death, and a recent study of yeast apoptosis recommend a combination of assays to determine the rate of yeast apoptosis [105]. Thus, further testing, preferably by a combination of assays to determine the cell viability, accumulation of RS, DNA fragmentation, and cell integrity, are required to measure the ratio of necrotic to apoptotic death in LEV-2 yeasts exposed to UV-B.

#### Other signalling proteins

Five protein kinases, not related to MAPK pathway, were identified in the yeast sample (**Error! Reference source not found.**). Of those, three adenylate kinases had low expression levels in samples subjected to 1 hour or longer UV irradiation, while two kinases of unknown specificity, exhibited significantly increased (fold change > 2) expression levels in yeast sample exposed to 24 hours of UVR. The protein kinases are involved in a large number of signalling pathways [116], and further research is required to elucidate the functions of these proteins in LEV-2.

### Proteins involved in biosynthesis of antioxidants

MudPIT analysis of yeast LEV-2 identified and quantified 17 proteins involved in biosynthesis of small-molecule antioxidants such as glutathione and ubiquinol. The identified proteins (**Error! Reference source not found.**) included enzymes involved in biosynthesis of glutathione [45] such as hydroxyacylglutathione hydrolases (HAGHs) and glutathione S-transferases (GSTs), enzymes involved in biosynthesis of vitamin B6 (PdxS/SNZ family lyases and pyridoxine 4-dehydrogenases) [46], and succinate dehydrogenase (SdHs) enzymes, which reduce oxidised coenzyme Q10 to its reduced form (ubiquinol) [47]. UV-mediated induction of oxidative stress and depletion of cellular small molecule antioxidants was previously observed in multiple models including animal cell lines, plants, yeasts and bacteria [3], [4]. For example, Heck et al. (2003) found that UV-B light is absorbed by catalase and increases the H_2_O_2_ generating activity of this enzyme in human and mouse keratinocytes, and in hamster fibroblasts [4]. A study by Podda et al. (1998) found that simulated solar UV radiation depletes small molecule antioxidants ubiquinol, α-tocopherol and ascorbic acid in the cell culture of human skin, in a linear, dose-dependent fashion [117]. UV radiation was shown also to induce RS generation in cyanobacteria [118], algae [119], and plants [120], and is also considered to be a major source of oxidative stress in yeasts [58]. In this study, the fold changes of the majority of enzymes involved in biosynthesis of small molecule antioxidants were moderately reduced in the LEV-2 yeast cultures irradiated for 1 hour to 8 hours (**Error! Reference source not found.**). This fold change reduction co-occurred with the reduction in antioxidant activity of cell extracts of LEV-2 yeast cultures exposed to 1 to 4 hours of UV-B, as measured by DPPH free radical quenching assay (**Error! Reference source not found.**), indicating that the UV-B radiation, and the associated oxidative stress, deplete the small molecule antioxidants of *Sporobolomyces* yeast LEV-2. These results are in agreement with the reports of Gasch et al. [60], [61], who used microarrays to study the protein expression of *S. cerevisiae* subjected to environmental stresses such as heat shock and exposure to H_2_O_2_ and found that temperature-induced stresses induced a fold change reduction in GST, SdHs and SNZ proteins [60], [61] of *S. cerevisiae*, consistent with the fold change patterns observed in this study. While the studies by Gasch et al. [60], [61] did not measure the effects of UV-B radiation, it has been shown previously that heat-shock, similarly to UV-B, induces oxidative stress in yeasts. That is because *S. cerevisiae* strains deficient in genes encoding antioxidant enzymes, such as catalase, SoD and cytochrome c peroxidase, are highly sensitive to heat-shock, while yeasts engineered to over-express these enzymes have an elevated resistance to heat shock and oxidative stress [121]. Notably, certain enzymes involved in biosynthesis of small-molecule antioxidants, such as glutamate cysteine ligase (GCL) and GSH synthetase (GS) [122], were not identified in this study. This is possibly due to the lack of the genome sequence for LEV-2 sample, which necessitated the use of related yeast proteomes for the database matching of MS spectra generated from LEV-2 proteins, and likely reduced the number of identified proteins.

### Enzymatic antioxidants

Enzymatic antioxidants include enzymes that convert RS into less reactive chemical species, such as superoxide dismutase (SOD), superoxide reductase (SOR) and catalase (CAT); enzymes involved in recycling of non-enzymatic small molecule antioxidants (e.g. glutathione reductase); and enzymatic systems that reduce oxidised cellular macromolecules, such as thioredoxin and glutaredoxin systems [123], [124]. While the individual enzymatic antioxidants are non-essential, presumably because cellular antioxidant systems have a considerable level of redundancy, the yeast *S. cerevisiae* [125], [126] and mouse animal models [127] deficient in certain antioxidant enzymes, such as mitochondrial SODs, were shown to be hypersensitive to oxidative stress. In addition, animal cell lines in which the biosynthesis of enzymatic antioxidants was induced by Nrf2 activators such as sulforaphane were found to be highly resistant to UV-induced oxidative stress [13] and to oxidants such as H_2_O_2_ [128]. Analogous increase in resistance to oxidative stress was also found in mouse models exposed to the Nrf2 activator sulforaphane [14], and in yeasts genetically engineered to over-express the bZip transcription factor Yap1 [129]. Enzymatic antioxidants are highly conserved across the domains of life, and the expression of human superoxide dismutase enzyme in yeast found was found to increase yeast resistance to oxidants such as paraquat [130].

Aldehyde dehydrogenases (ADHs) are involved in the metabolism of toxic aldehydes produced during oxidative stress [49]. One ADH and a benzaldehyde dehydrogenase were identified and quantified by the MudPIT analysis of *Sporobolomyces* yeast LEV-2, and both enzymes showed fold change increase in LEV-2 cultures exposed to 24 hours of UV-B irradiation (**Error! Reference source not found.**). Superoxide dismutases (SODs) catalyse the conversion of a superoxide anion to H_2_O_2_ and O_2_, and play a critical role in protection against oxidative stress in all eukaryotes, including yeasts [48]. Four SODs were identified in this study, and the expression levels of SODs were increased in UV-B irradiated LEV-2 cultures (**Error! Reference source not found.**). This fold change increase of ADHs and SODs correlated with the increase in antioxidant, DPPH quenching, activity of extracts of yeast LEV-2 cultures exposed to 8 hours and 24 hours of UV-B (**Error! Reference source not found.**A), and with the observed reduction in yeast LEV-2 cell death (**Error! Reference source not found.**), indicating that these enzymes protect yeast LEV-2 against UV-induced oxidative stress. These results are consistent with previous studies of yeast *S. cerevisiae* exposed to oxidants such as H_2_O_2_ and to heat-shock induced oxidative stress [61], and suggests that the function of SODs and ADHs is conserved between *S. cerevisiae* and the yeast LEV-2.

Catalases (Cat) facilitate the breakdown of H_2_O_2_ to O_2_ and H_2_O, and are major antioxidant enzymes in yeasts [48]. A catalase identified in this study showed a moderate fold change reduction in yeast cultures irradiated for 2 hours to 8 hours ((**Error! Reference source not found.**). Cytochrome C peroxidases (CCPs) catalyse the conversion of H_2_O_2_ to H_2_O, and have been implicated in yeast response to heat shock and oxidative stress [48], [50]. All of the five identified CCPs had reduced expression levels in yeast samples exposed to moderate duration of UV-B doses (1 hour to 4 hours), and the results were ambiguous for LEV-2 cultures irradaated for 8 hours and 24 hours. Isocitrate dehydrogenase (IDH) enzymes are involved in citric acid cycle, and catalyse the two-step oxidative carboxylation of isocitrate to α-ketoglutarate. IDHs exist in multiple isoforms which differ in the use of NAD+ or NADP+ as a cofactor, and in cellular localization to cytosol or mitochondria. While not direct antioxidants, IDHs play a part in cellular defences against RS by reducing NAD(P)+ to NAD(P)H to enable the regeneration of glutathione (GHS) and thioredoxins [52], [53]. This study identified two NAD+ dependant IDHs, both of which showed fold change increase in LEV-2 cultures irradiated for 8 hours and 24 hours, and four NADP+ associated IDHs which displayed fold change reduction after 1 hour or longer UV-B exposure. The observed fold change reduction of LEV-2 IDH, CCP and Cat enzymes corresponds to the reduction in DPPH quenching observed for LEV-2 cultures exposed to 1 hour to 4 hours of UV-B (**Error! Reference source not found.**), and is possibly a result of depletion of these enzymes by UV-B induced oxidative stress. Oxidative stress was reported to deplete antioxidants, especially GSH, in the yeast *S. cerevisiae* [131], but the extent of antioxidant depletion varied with the source and duration of stress [48], [61]. The depletion of GSH was reported also for UV-irradiated human skin models [117], indicating that oxidative stress has a similar effect on enzymatic antioxidants in diverse eukaryotic organisms.

Glutathione peroxidase (Gpx) enzymes catalyse conversion of RS, such as H_2_O_2_, to non-reactive compounds such as H_2_O, and are major eukaryotic enzymatic antioxidants [51]. Two Gpx enzymes were identified in this study, and the lack of major fold changes of these enzymes (**Error! Reference source not found.**) indicates that Gpx enzymes are stable during the oxidative stress. This is possibly because these enzymes catalyse RS conversion (ROOH + 2GSH à ROH + GSSG + H_2_O reaction) rather than the direct reduction of RS. This is in agreement with previous yeast studies of Gasch et al. (2000) and Yoshimoto et al. (2002), who identified that Gpx-encoding mRNAs do not exhibit significant changes in stressed yeast *S. cerevisiae* (majority of identified fold-changes were <= 1.5) [61], [83].

### DNA repair and replication

UV radiation is a genotoxic environmental agent that inflicts DNA damage by causing the oxidative stress and by inflicting direct damage to the DNA macromolecule. UV-A and UV-B light induce the cellular production of reactive oxygen-derived species, such as H_2_O_2_ [3], [4], and the evidence for UV-induced oxidative damage has been found in algae [119], plants [120], animals [4], [117] and bacteria [57]. UV-induced oxidative stress is also considered to be a major source of UV-induced damage in yeasts [132]. During oxidative stress, the highly reactive hydroxyl radicals (OH•) react with the DNA molecule and cause the formation of single and double stranded DNA breaks, base modifications and cross-linkage with proteins [133]. In addition to inducing oxidative stress, UV-B light is also absorbed by DNA pyrimidine bases, thymine and cytosine, and induces photoreactions that lead to formation of mutagenic DNA photoproducts, such as cyclobutane–pyrimidine dimers (CPDs) and 6–4 photoproducts [134], as well as formation of single stranded DNA breaks [2].

The maintenance of DNA is essential for all forms of life, and DNA repair mechanisms are well conserved across all domains of life, including bacteria, yeasts and animals [135]. The photoreactivation mechanism that utilizes photolyase enzymes to repair CPDs and 6–4 photoproducts is considered to be the oldest and the simplest mechanism of DNA repair, as evinced by existence of photolyases in all domains of life, including archaea, and by the fact that it only utilizes a single enzyme [136]. Contrasted to photoreactivation are excision repair mechanisms that do not repair the DNA damage, but instead remove the damaged section of DNA and replace it with newly synthesized nucleotides [132]. These mechanisms comprise three major categories: base excision repair (BER), which repairs small changes in DNA that do not alter the DNA helix structure; nucleotide excision repair (NER) which replaces “bulky” DNA adducts such as thymine dimers and 6,4-photoproducts; and mismatch repair mechanisms that repair erroneous insertions, deletions and incorporations of nucleic bases during DNA replication and recombination [135], [137]. Unlike the photoreactivation which utilizes only a single enzyme, excision repair pathways are comprised of numerous enzymes. For example, the BER pathway utilizes DNA glycosylase to recognize DNA damage and remove damaged nucleic base; apurinic/apyrimidinic (AP) endonuclease, AP lyase and phosphodiesterase to excise the deoxyribose phosphate residue left-over after removal of damaged nucleic base; DNA polymerase to repair the introduced DNA gap; and DNA ligase to connect the newly synthesized DNA with the rest of the DNA strand. The NER mechanism is also comprised of multiple enzymes including proteins that recognize DNA damage, endonucleases, helicases, ligases and other proteins required for regulation of the process [132]. Notably, not all organisms possess all of the described DNA repair mechanisms; for example, photolyases required for photoreactivation repair of CPDs have been reported in bacteria, fungi, plants, invertebrates and many vertebrates, but not in humans, while photolyases that reverse 6–4 photoproducts have been found in fly *Drosophila*, frog *Xenopus laevis*, and certain snakes, but not in *E. coli*, yeast *S. cerevisiae* or humans [132].

Six yeast enzymes involved in the repair and replication of DNA were identified and quantified in this study (**Error! Reference source not found.**). Identified proteins included DNA polymerases, DNA helicase, exonuclease and DNA ligase, all of which are involved in repair of single strand DNA breaks by BER and in the repair of double-strand breaks by non-homologous end-joining [138], [139]. In addition, one dUTP pyrophosphatase enzyme was identified and quantified; this enzyme is essential for DNA replication and repair as it prevents erroneous incorporation of uracil into the DNA [140]. The expression levels of these enzymes were increased in the yeast LEV-2 sample exposed to 24 hours of UV irradiation (**Error! Reference source not found.**), indicating that long-term UV-induced stress activates DNA repair in yeast LEV-2. This increase in expression of DNA repair enzymes correlated with the reduction in yeast cell death (**Error! Reference source not found.**), suggesting that the yeast LEV-2 adapts to UV-induced stress by increasing the rate of DNA repair. Notably, photolyase enzymes were not identified in this study, possibly because these proteins have high molecular mass and charge (for example, the yeast *S. cerevisiae* photolyase encoded by *PHR1* gene is a 66 kDa protein with pI value of 9.2), and it was demonstrated that MudPIT detection rates are low for large, charged proteins [141].

### Heat-shock proteins and related proteins

Heat shock proteins (HSPs) and chaperones are highly conserved across all domains of life [142]. HSPs are mainly involved in resistance to heat-induced stress, while chaperones also play a role in protein folding and unfolding, assembly of protein complexes, protein transport, cell-cycle control and protection against apoptosis [143]. The MudPIT analysis of LEV-2 yeast exposed to long-term UV irradiation identified and quantified 24 proteins annotated as heat-shock proteins or chaperones (**Error! Reference source not found.**). The expression of the majority of HSPs was increased in samples exposed to 24 hours of UV-B, co-occurring with the increase in expression of other stress resistance proteins (**Error! Reference source not found.**) and reduction in cell death (**Error! Reference source not found.**). This increase in expression was particularly noticeable for 10 kDa mitochondrial HSPs, Chaperonin ATPases TCP-1 and 90 kDa HSP proteins, indicating these families of HSP proteins play a major part of stress response in yeast LEV-2. This is possibly because the 90 kDa HSPs are involved in cellular signalling in addition to the function in stress response [144], and the TCP-1 chaperonins also mediate the ATP-dependent renaturation of proteins to assist in repair of UV and oxidative damage [145]. Interestingly, the expression levels of multiple HSP proteins, classified as 60 kDa HSPs and 70 kDa HSPs, were reduced in the LEV-2 samples exposed to 1 hour - 4 hours of UV-B (**Error! Reference source not found.**). These results match a previous study of Gasch et al. (2000) that measured mRNA levels in yeast *S. cerevisiae* exposed to different sources of stress, and identified a significant increase in levels of mRNAs encoding 12 kDa, 30 kDa, 42 kDa and 104 kDa HSPs during heat shock, while levels of mRNAs encoding other HSPs were largely unchanged [61]. The related study of Yoshimoto et al. (2002), measured mRNA levels of yeast *S. cerevisiae* under different environmental stresses (not including UV), and also identified that changes in levels of HSP-encoding mRNAs vary with the source and the duration of stress [61], [83]. The reduction in HSP levels observed for UV-exposed yeast LEV-2 is possibly due to the reduction in protein biosynthesis observed in samples exposed to 1 hour, 2 hours and 4 hours of UV-B (**Error! Reference source not found.**) and indicates that 60 kDa HSPs and 70 kDa HSPs are unlikely to be a major part of UV response in this yeast.

### Proposed model of yeast LEV-2 stress response

The results of the quantified proteome of LEV-2 exposed to extended UV-B irradiation, were related to the antioxidant activity of cell-lysis extracts of LEV-2 cultures, measured by the DPPH assay (**Error! Reference source not found.**), and to survival rates of LEV-2 cultures exposed to long-term UV-B irradiation (**Error! Reference source not found.**) to construct the model of UV-stress response of the *Sporobolomyces* yeast LEV-2. This comparison of yeast viability, antioxidant activity and protein fold changes of UV-exposed LEV-2 suggested that a moderately long UV-B irradiation (∼1 hour) caused the fold change reduction in the yeast LEV-2 proteins involved in the carbohydrate metabolism, energy metabolism and protein biosynthesis (**Error! Reference source not found.**), caused the moderate (∼25%) reduction in the yeast viability (**Error! Reference source not found.**), and triggered fold change increase in cellular signalling proteins related to Ras and Yap1 pathways (**Error! Reference source not found.**, **Error! Reference source not found.**). The continued irradiation further reduced the energy metabolism and viability of yeast cultures, with the maximal impact observed after 4 hours of UV-B. The expression level fold changes of proteins involved in yeast energy metabolism, however, increased after 8 hours of UV-B exposure and reverted to levels close to the control after 24 hours of UV-B exposure (**Error! Reference source not found.**). This co-occurred with the reduction of yeast cell death, observed after 8 hours or longer UV-B exposure (**Error! Reference source not found.**), and with an increase in expression levels of LEV-2 stress-response proteins such as enzymatic antioxidants (**Error! Reference source not found.**) and certain heat-shock proteins (**Error! Reference source not found.**). The levels of stress-response proteins also correlated with the free-radical quenching activity of the yeast cell extracts, as measured by the DPPH free radical scavenging assay (**Error! Reference source not found.**). These results can be best explained by proposing a yeast stress response and adaptation model, illustrated in the Figure 11. The model postulates that yeast UV response can be modelled in a four-step process: 1) UV exposure inflicts direct damage to the cell and induces oxidative stress, which in the short term induces bZip protein mediated signalling, analogous to Nrf2 signalling in animals or Yap1 signalling in the yeast *S. cerevisiae* (“signalling” phase); 2) The damage caused by the extended exposure to UV impairs carbohydrate and energy metabolism of the cell, and RS deplete cellular antioxidants (“stress” phase); 3) As the cellular defences are expressed, the yeast undergoes an “adaptation” phase where newly synthetized antioxidant enzymes (SODs, Cat, GTRx and NQR) and small molecule antioxidants (e.g. glutathione, ascorbic acid and α-tocopherol), DNA-repair enzymes, and UV-protective metabolites such as UV-absorbing carotenoids reduce the stress level and cellular metabolism starts recovery; 4) this leads to a “stress resistant” phase of growth where cellular metabolism and growth stabilize at a rate lower then pre-stress conditions.

**Figure 11:**
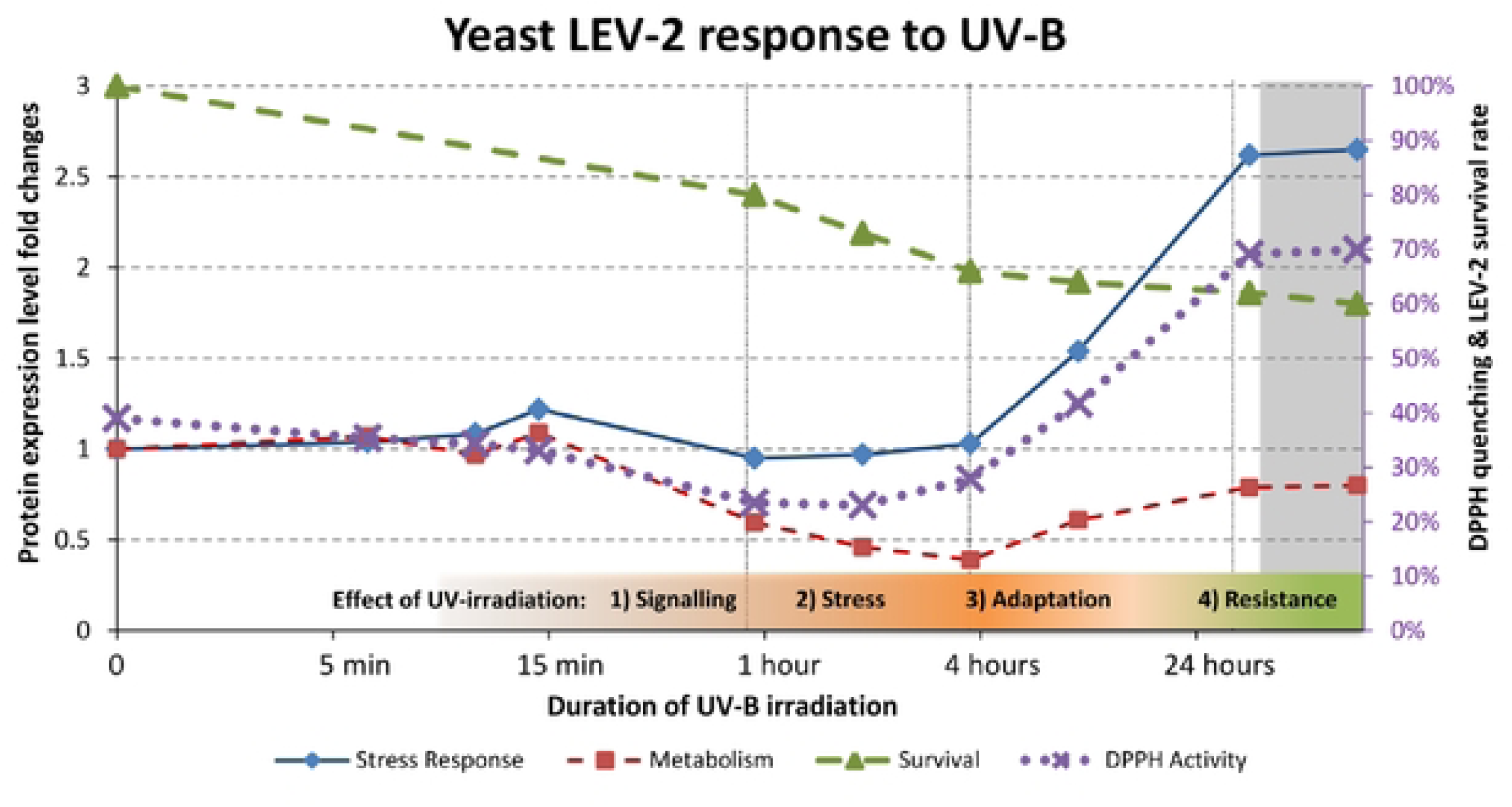
Proposed proteomics-based cellular response model for the *Sporobolomyces* yeast LEV-2 response to extended UV-8 exposure. The proposed model of stress response of UV-tolerant yeast LEV-2. The fold changes of proteins involved in stress response (blue line), and proteins involved in carbohydrate metabolism (red line) correspond to main (left side) Y-axis. The yeast survival curve (green line) and the DPPH quenching activity of yeast cell extracts (purple line) are expressed in percentages and correspond to the secondary (right side) Y-axis. The phases of stress response are denoted on the bottom on the chart The chart is based on MudPIT experiment of LEV-2 yeast, and the grey shaded part of the chart represents the predicted patterns of yeast proteome, based on the “adaptation model” (see text for details).

The induction of stress resistance by low-intensity, sub-lethal, stress has been previously described in different eukaryotic models, including the yeast *S. cerevisiae* [146], mammalian cell cultures [147] and *in-vivo* animal models [14]. For example, in a study by Davies et al. (1995), *S. cerevisiae* cultures conditioned by exposure to low concentration (0.4 mM) of H_2_O_2_ were found to survive, with ∼90% viability rate, the subsequent exposure to high concentration (3 mM) of H_2_O_2_, lethal to unconditioned yeasts. The microarray studies of *S. cerevisiae* exposed to different environmental stresses, such as heat-shock, H_2_O_2_ and toxic metals, found also that induction of proteins involved in stress response is transient, and the expression levels revert to levels close to non-stressed yeast during the prolonged stress as yeast adapts to stress [60]. Similar adaptation was observed for the fly *D. melanogaster* and for mouse cell cultures, where the exposure to a low dose of H_2_O_2_ conditioned the observed animals or cells to the following oxidative shock caused by high concentration of H_2_O_2_ [148]. In animals, the adaptation to oxidative stress is mediated by the Nrf2 pathway; The low dose of oxidant, such as H_2_O_2_, induces the transcription of antioxidant and cytoprotective genes regulated by the bZip transcription factor Nrf2, and stimulates an increase in tolerance to oxidative stress [5]. Numerous studies of mouse models and animal cell lines have demonstrated that pre-treatment by Nrf2-activators such as SFN increased tolerance to oxidative stress in animals [6]. For example, Nrf2 upregulation was found to protect human cells against cigarette smoke [149], UV-induced oxidative damage [150] and toxicity of drugs such as cisplatin [11]. The activation of Nrf2 was also shown to protect mouse models against UV-induced carcinogenesis [14], while the Nrf2-knockout mouse models were shown to lack the ability to adapt to oxidative stress caused by carcinogens such as benzo[a]pyrene [66], and by drugs such as acetaminophen [67]. bZip transcription factors have been shown to be evolutionary conserved between yeasts and animals [20], [21], and *S. cerevisiae* bZip protein Yap1 regulates the response to oxidative stress in yeast [26], indicating that the molecular mechanism of adaptation to oxidative stress is evolutionary conserved between yeasts and animals.

In summary, in this study the *Sporobolomyces* yeast LEV-2 bZip protein LEV-2_XP_007274754.1 showed a significant fold change increase in yeast cultures exposed to UV-B (**Error! Reference source not found.**). The increase in expression of the bZip protein matched the results of previous studies that showed increase in expression of *S. cerevisiae* Yap1 protein in yeast exposed to oxidative stress [76] and the increase in bZip protein in animal cells exposed to oxidants [78], [79]. This suggests that the LEV-2 bZip protein LEV-2_XP_007274754.1 is a homolog of the *S. cerevisiae* transcription factor Yap1 and the vertebrate transcription factor Nrf2, and that the bZip protein mediated response to oxidative stress is conserved between LEV-2, *S. cerevisiae* and animals, as was previously predicted by our *in-silico* studies [20], [21]. It should be noted, however, that further studies are required to quantify the relative contributions of different signalling pathways in stress response of yeast LEV-2.

## Conclusions

The primary objective of this study was to determine if homologs of the vertebrate bZip protein Nrf2 play a role in stress response of UV-tolerant yeasts, such as carotenoid-producing yeasts of Genus *Sporobolomyces.* The quantitative MudPIT analysis of the *Sporobolomyces* yeast LEV-2 exposed to long term UV-B irradiation was followed by the functional annotation of the yeast LEV-2 proteome. This study identified four basic leucine zipper (bZip) proteins and suggested that a bZip protein designated LEV-2_XP_007274754.1 is a homolog of the yeast bZip protein Yap1 and animal bZip protein Nrf2, and initiates the response to UV-induced oxidative stress in yeast LEV-2. The secondary goal of this study was to describe the UV-stress response of yeast LEV-2, and to evaluate if the stress response is conserved between LEV-2, other yeasts such as *S. cerevisiae* and animal models. A quantitative MudPIT proteomics analysis of LEV-2 cultures exposed to extended UV-B irradiation led to the proposal of a 4-step model, where 1) short-term UV irradiation induces cellular signalling; 2) depletion of cellular antioxidants and reduction in cellular metabolism; 3) adaptation by high expression of antioxidant enzymes and production of UV-absorbing red pigments; and 4) shift into stress-resistant phase of growth characterized by high expression of antioxidants and reduced metabolism. This model (**Error! Reference source not found.**) matches the adaptation to stress observed also in *S. cerevisiae* and in animals, indicating that the molecular mechanisms of adaptation to oxidative stress are evolutionary conserved between *Basidiomycota* fungi, *Ascomycota* fungi, and animals.

## Acknowledgements

We thank Dr Howard G Wildman from Microbial Management Systems Australia for his advice on yeast isolation during the early stages of this work. This work was supported by the United Kingdom Medical Research Council (MRC Doctoral Training Program Grant G82144A).

## Notes

### Competing Interest Statement

The authors have declared no competing interest.

